# Life history recorded in the vagino-cervical microbiome

**DOI:** 10.1101/533588

**Authors:** Zhuye Jie, Chen Chen, Lilan Hao, Fei Li, Liju Song, Xiaowei Zhang, Jie Zhu, Liu Tian, Xin Tong, Kaiye Cai, Yanmei Ju, Xinlei Yu, Ying Li, Hongcheng Zhou, Haorong Lu, Xuemei Qiu, Qiang Li, Yunli Liao, Dongsheng Zhou, Heng Lian, Yong Zuo, Xiaomin Chen, Weiqiao Rao, Yan Ren, Yuan Wang, Jin Zi, Rong Wang, Na Liu, Jinghua Wu, Wei Zhang, Xiao Liu, Yang Zong, Weibin Liu, Liang Xiao, Yong Hou, Xun Xu, Huanming Yang, Jian Wang, Karsten Kristiansen, Huijue Jia

**Affiliations:** BGI-Shenzhen, Shenzhen 518083, China; Shenzhen Key Laboratory of Human Commensal Microorganisms and Health Research, BGI-Shenzhen, Shenzhen, China; Department of Biology, Ole Maaløes Vej 5, University of Copenhagen, Copenhagen, Denmark; Shenzhen Engineering Laboratory of Detection and Intervention of human intestinal microbiome, BGI-Shenzhen, Shenzhen, China; BGI Education Center, University of Chinese Academy of Sciences, BGI-Shenzhen, Shenzhen 518083, China; China National Genebank, BGI-Shenzhen, Shenzhen 518120, China; BGI-Qingdao, BGI-Shenzhen, Qingdao, 266555, China; James D. Watson Institute of Genome Sciences, Hangzhou, China

## Abstract

The vagina contains at least a billion microbial cells, which are dominated by Lactobacilli. Here we perform metagenomic shotgun sequencing on cervical samples from 1148 women. Factors such as pregnancy, delivery histories and breast-feeding were all more important than menstrual cycle in shaping the microbiome. *Bifidobacterium breve* was seen with older age at sexual debut; *Lactobacillus crispatus* negatively correlated with pregnancy history; potential markers for lack of menstrual regularity, heavy flow, dysmenorrhea, contraceptives were also identified. Other features such as mood fluctuations and facial speckles could potentially be deduced from the vagino-cervical microbiome. Gut and oral microbiome, plasma vitamins, metals, amino acids and hormones showed associations with the vagino-cervical microbiome. Our results offer an unprecedented glimpse into the microbiota of the female reproductive tract and call for international collaborations to better understand its long-term health impact.

**Highlights:** Shotgun sequencing of 1148 vagino-cervical samples reveal subtypes;

Other omics such as moods and skin features could be predicted by the vagino-cervical bacteria;

Factors such as delivery mode and breast-feeding associate with the vagino-cervical microbiome;

With dwindled Lactobacilli, postmenopausal samples are relatively enriched for viruses.

## INTRODUCTION

The human body is a supra-organism containing tens of trillions of microbial cells (Sender et al., 2016). Studies in human cohorts and animal models have revealed an integral role played by the gut microbiota in metabolic and immunological functions (Sommer and Bäckhed, 2013). Disease markers as well as prebiotic or probiotic interventions are being actively developed (O’Toole et al., 2017; Wang and Jia, 2016). Studies on other mucosal sites such as the vagina and the mouth lag behind (Lloyd-Price et al., 2017; The Human Microbiome Project Consortium., 2012; Zhang et al., 2015b); the potential for the microbiota in these body sites has not been fully realized. The presence of microorganisms beyond the cervix (i.e. the upper reproductive tract) is increasingly recognized even in non-infectious conditions (Aagaard et al., 2014; Chen et al., 2017; Li et al., 2018), with much debated implications for women’s and infants’ health.

Lactobacilli have long been regarded as the keystone species of the vaginal microbiota. Lactic acid produced by these microorganisms helps maintain a low vaginal pH of 3.5-4.5, and wards off pathogenic micro-organisms (Ma et al., 2012). Prevention of the human immunodeficiency virus (HIV) and other sexually transmitted infections (STIs), preterm birth and bacterial vaginosis (BV) has been a major effort. Germ-free mice treated with *L. crispatus* had fewer activated CD4+ T cells in the genital tract compared to those treated with *Prevotella bivia*, explaining the difference in HIV acquisition in human cohorts (Gosmann et al., 2017). How vaginal Lactobacilli interplay with *Candida albicans* and other related fungi is also an important question for preventing or treating vulvovaginal candidiasis (Bradford and Ravel, 2017). The over 90% of human sequences in female reproductive tract samples, in contrast to 1% in feces (Li et al., 2018; Methé et al., 2012; Wang and Jia, 2016), has made metagenomic shotgun sequencing more expensive. Most studies of the vaginal microbiota used 16S rRNA gene amplicon sequencing, which lacked a view of the overall microorganism communities including bacteria, archaea, viruses and fungi (Byrd et al., 2018), as well as the functional capacity encoded. Besides infection, studies on current sexual activity and the menstrual cycle occurred naturally to the vaginal microbiota field (Gajer et al., 2012; Ravel et al., 2010). However, lasting impacts from other potentially important factors such as sexual debut, pregnancy and breast-feeding have not been looked at in a reasonably large cohort.

As a sizable reservoir of microbes instead of a transient entity, the female reproductive tract microbiota might also reflect conditions in other body sites. It is however not clear whether other omics in circulation or in the intestine cross-talks with the vagino-cervical microbiome. Intersecting with hormones, metabolic and immune functions, we find it intriguing to explore the potential link of the vagino-cervical microbiome to the brain and the face.

Here, we report the full spectrum of the vagino-cervical microbiome from 1148 healthy women using metagenomic shotgun sequencing. As part of the 4D-SZ (trans-omic, with more time points in future studies, based in China) cohort, we also comprehensively measured parameters such as fecal/oral microbiome, plasma metabolites, medical test data, immune indices, physical fitness test, facial skin imaging, as well as female life history questionnaire, lifestyle questionnaire, and psychological questionnaire (Figure 1). Our work pinpoints other metadata or omics that can predict or be predicted from the microbiota in the female reproductive tract, which would illuminate future designs of population cohorts, mechanistic investigations, as well as means of intervention.

**Figure 1.**
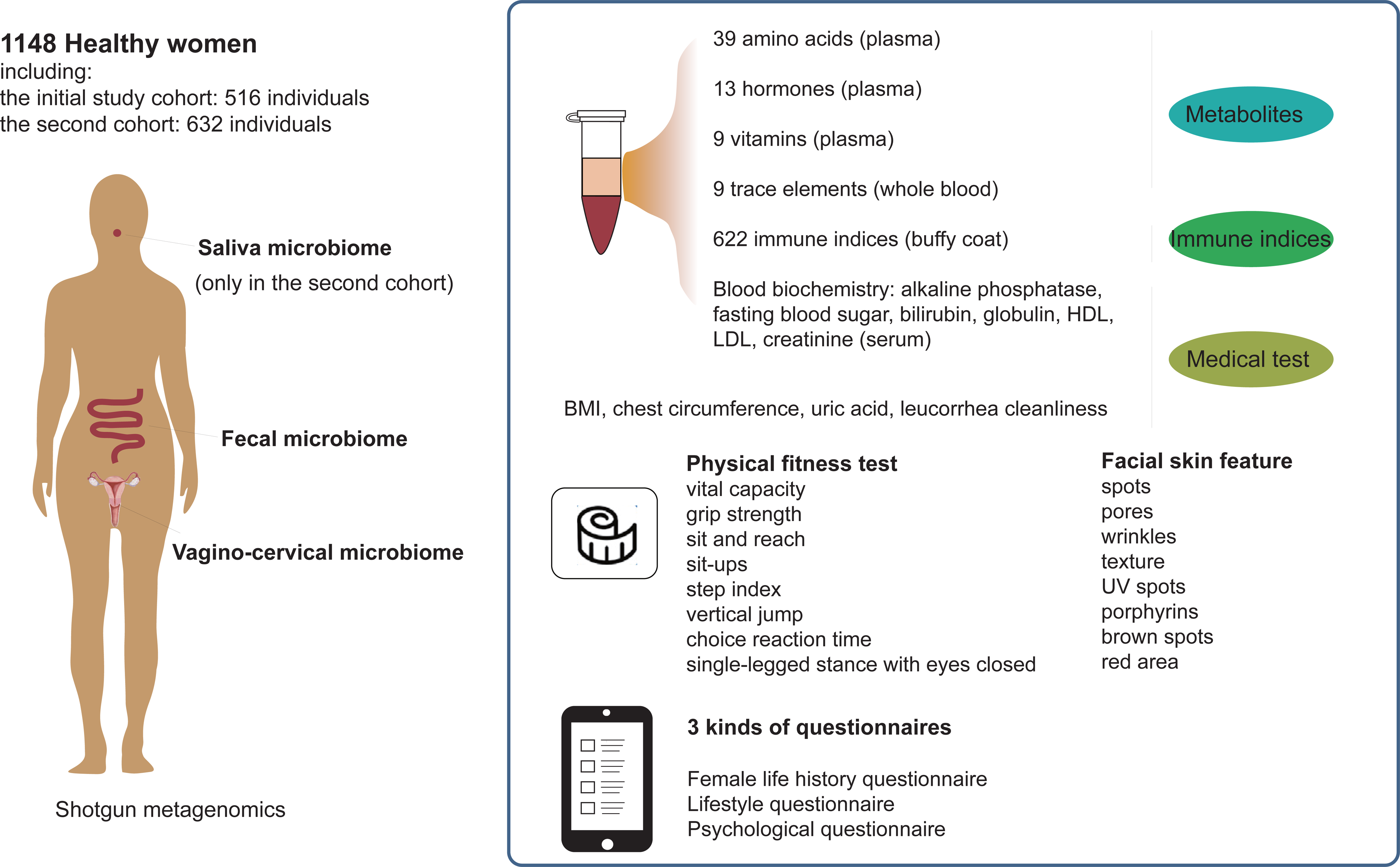
Schematic overview of the study. The study consists two cohorts with varying characteristics. 516 volunteers recruited in 2017 are regarded as the initial study cohort. A second cohort containing 632 volunteers were recruited in 2018. Vagino-cervical microbiome, fecal microbiome, metabolites, medical test data, physical fitness test data, facial skin feature and three kinds of questionnaires are collected in both cohorts. Saliva microbiome is only available in the second cohort and the woman life questionnaires differ by a few terms.

## RESULTS

### Dominant bacterial and non-bacterial members of the vagino-cervical metagenome

To explore the vagino-cervical microbiome, 516 healthy Chinese women aged 21-52 (median 29, 95% CI: 23–39) were recruited during a physical examination as the initial study cohort (Figure 1; Table S1A). Metagenomic shotgun sequencing was performed on the cervical samples, and high-quality non-human reads were used for taxonomic profiling of the vagino-cervical microbiome (Figure 2A; Table S1B).

**Figure 2.**
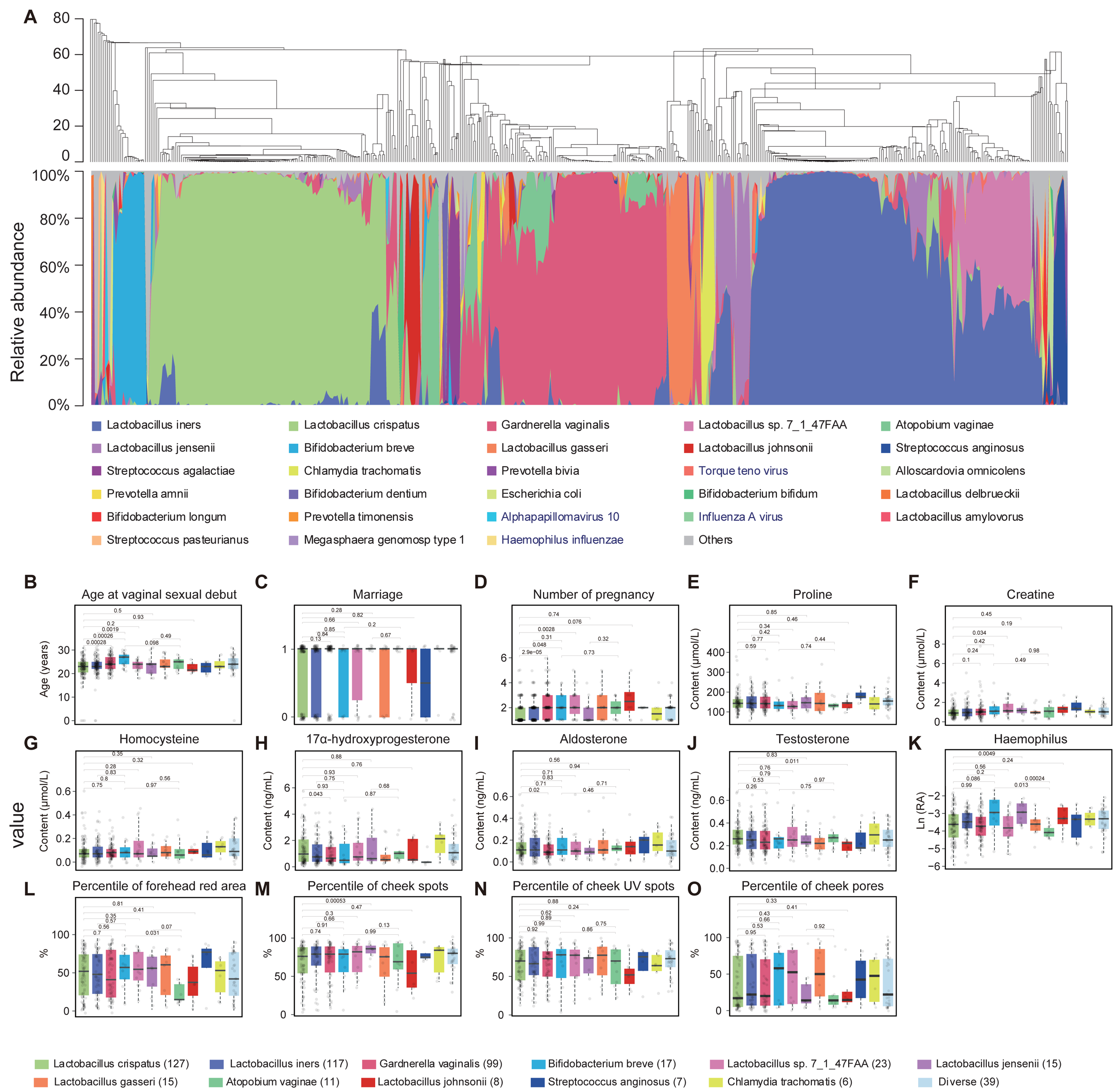
Vagino-cervical microbiome of the initial study cohort. (A) The microbial composition in each sample at the species level according to MetaPhlAn2 is shown. The samples were hierarchically clustered (R base hcluster function with centroid linkage based on Euclidean distance. Black and blue taxa labels used in legend denote bacteria and viruses, respectively. (B-O) Specific omics factors that shows significant difference among 14 key vaginal microbiota types estimated by GLM likelihood ratio test (p<0.05, 554 comparisons). T test use for post hoc compare. RA stands for relative abundance. The genus in sub-figure (M), (N), (O), (P) belongs to fecal microbiome. Boxes denote the interquartile range (IQR) between the first and third quartiles (25th and 75th percentiles, respectively), and the line inside the boxes denote the median. The whiskers denote the lowest and highest values within 1.5 times the IQR from the first and third quartiles, respectively.

In agreement with 16S rRNA gene amplicon sequencing data from the US (Ravel et al., 2010), the vagino-cervical microbiota of this Asian cohort was mostly Lactobacilli-dominated, while 19.19% of women harbored over 50% *Gardnerella vaginalis* (Figure 2A, Figure S1A). Although *A. vaginae* and *G. vaginalis* were commonly believed to co-occur in bacterial vaginosis (BV) (Fredricks et al., 2005; Serrano et al., 2019), 4 of the 10 volunteers with >50% in their cervical samples appeared to have no *G. vaginalis* (Figure 2A). The groups showed different characteristics in other omics (Figure 2B-O, Generalized linear model (GLM), p <0.05). 22.67% of the cohort was dominated by *Lactobacillus iners* (Figure 2A, Figure S1A), which was far less protective against bacterial and viral infections compared to *Lactobacillus crispatus* (Gosmann et al., 2017; Petricevic et al., 2014). The type characterized by *L. crispatus* was overrepresented in the women who had fewer pregnancies (Figure 2C, 2D). The plasma concentrations of 17 -hydroxyprogesterone and aldosterone were higher in individuals of the *L. crispatus*-α type than in *G. vaginalis* type individuals (Figure 2H-2J, p <0.05). Rare subtypes (> 50% relative abundance in less than 5% of the individuals) such as *Bifidobacterium breve* (3.29%)*, Lactobacillus jensenii* (2.91%)*, Lactobacillus gasseri* (2.91%)*, Atopobium vaginae* (2.31%)*, Lactobacillus johnsonii* (1.55%)*, Streptococcus anginosis* (1.36%)*, and Clamydia trachomatis* (1.16%) were also detected in this cohort (Figure 2A, Figure S1A). Individuals dominated by *B. breve* had older age at vaginal sexual debut compared to those dominated by *L. cripspatus* or *L. iners* (Figure 2B, p = 0.00026 and p = 0.0019, respectively). *Streptococcus agalactiae* (Group B *Streptococcus*), a bacterium responsible for neonatal sepsis and recently reported in placenta (de Goffau et al., 2019), could be detected in 5.62 % of the individuals (Figure 2A, Table S1C). Other microorganisms including *Prevotella bivia*, *Escherichia coli*, *Ureaplasma parvum*, human papillomavirus (HPV), herpesviruses, Influenza A virus and *Haemophilus influenzae* were abundant in some individuals (Figure 2A, Table S1C). According to the metagenomic data, the mean proportion of non-bacterial sequences was 3.45% (Figure 2A). The vaginal types (> 50% relative abundance for the single bacterium) of *G. vaginalis*, *L. crispatus*, and *B. breve* showed lower proportions of human sequences than other types (e.g. p = 3.9e-6 between *L. crispatus* and *L. iners* types, p = 1.7e-8 between *G. vaginalis* and *L. iners* types, p = 0.0036 between *B. breve* and *L. gasseri,* Wilcoxon ranked sum test, Figure S1B), suggesting that in future studies lower amounts of sequencing could be used for these compared to other types.

### Factors shaping the vagino-cervical microbiome and beyond

We computed the prediction value (5-fold cross-validated random forest model) of each factor in the female life history questionnaire on the microbiome data independently, and found the most important factors to be pregnancy history, marriage, number of pregnancies, number of vaginal deliveries, age at vaginal sexual debut, followed by age of marriage, current breast-feeding, and number of caesarean sections (Figure 3, 999 permutations and Benjamini-Hochberg tests, q < 0.05). Number of pregnancies correlated with pregnancy history (Spearman’s correlation coefficient (cc) = 0.916) and with number of abortions (Spearman’s cc = 0.699); Age at first pregnancy correlated age at most recent pregnancy (Spearman’s cc = 0.636).

**Figure 3.**
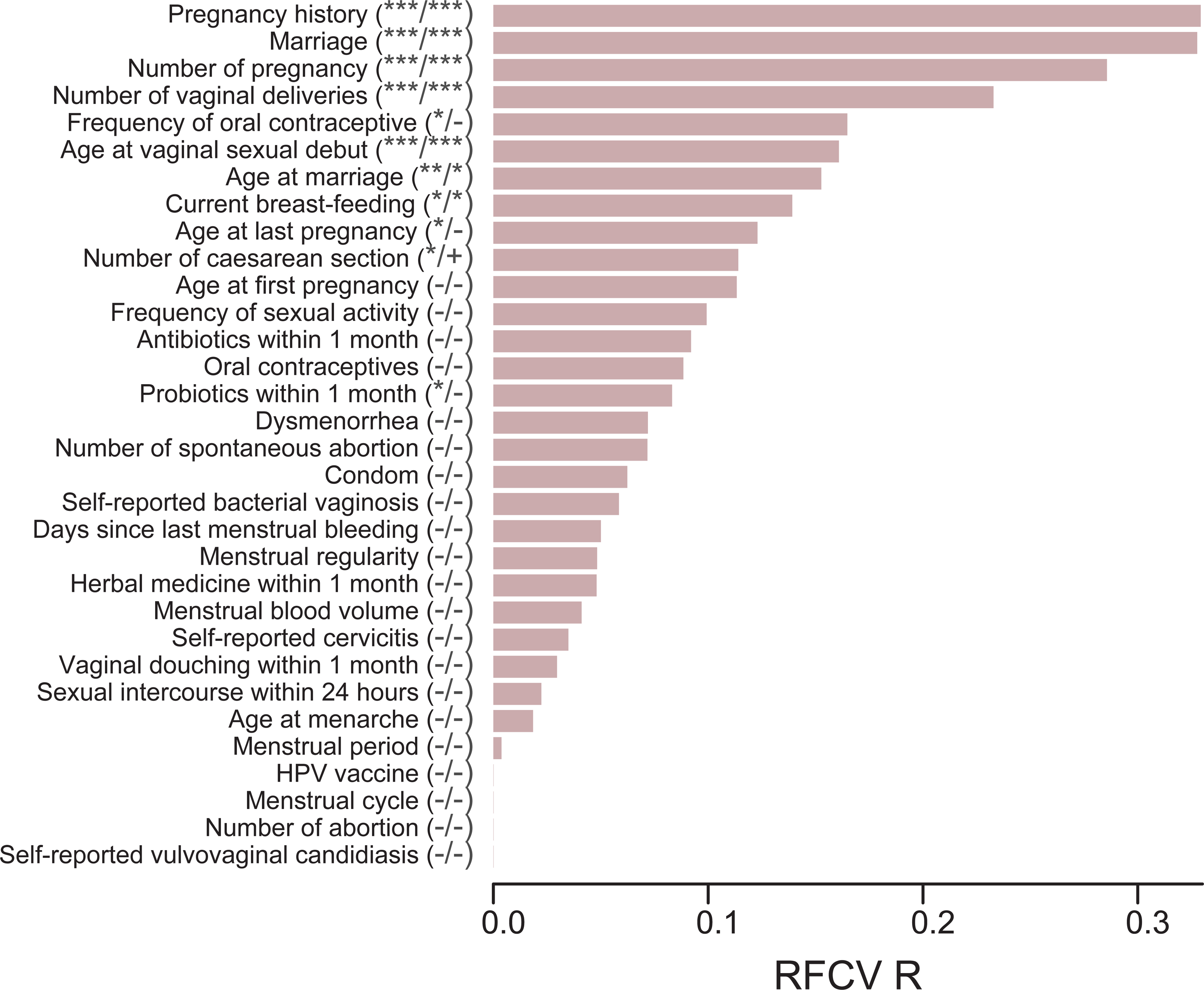
Factors from female life history questionnaire influencing the vagino-cervical microbiome in the initial study cohort. Female life history questionnaire entries on the vagino-cervical microbiome, ordered according to their 5-fold cross-validated random forest (RFCV) importance on the microbiome composition. X-axis (length of the bar) is the model performance measured as the spearman’s correlation between the prediction and measurement. First column stars after y-axis label are 999 times permutation p-value, second column stars are BH adjust p-value (32 comparisons), an “+” denotes Q-value <0.1, an asterisk denotes Q-value <0.05, two asterisks denote Q-value <0.01, three asterisks denote Q-value <0.001.

These important factors were validated in an independent cohort of 632 individuals that differed in age distribution as well as sequencing mode (Figure 1, Figure S2, Table S1E, paired-end 100bp for vagino-cervical samples). Questionnaire entries such as pregnancy history, marital status, current breast-feeding, and mode of the most recent delivery were again found as the most important factors to influence the vagino-cervical microbiome (Figure S2). The strong signal with duration of current breast-feeding reflected difference in questionnaire design that was only available in one but not the other cohort (Figure 4, Table S1A, Table S1E), while the impact from menstrual cycle was partly augmented with the presence of postmenopausal women in the second cohort (Figure 3, Figure S2).

**Figure 4.**
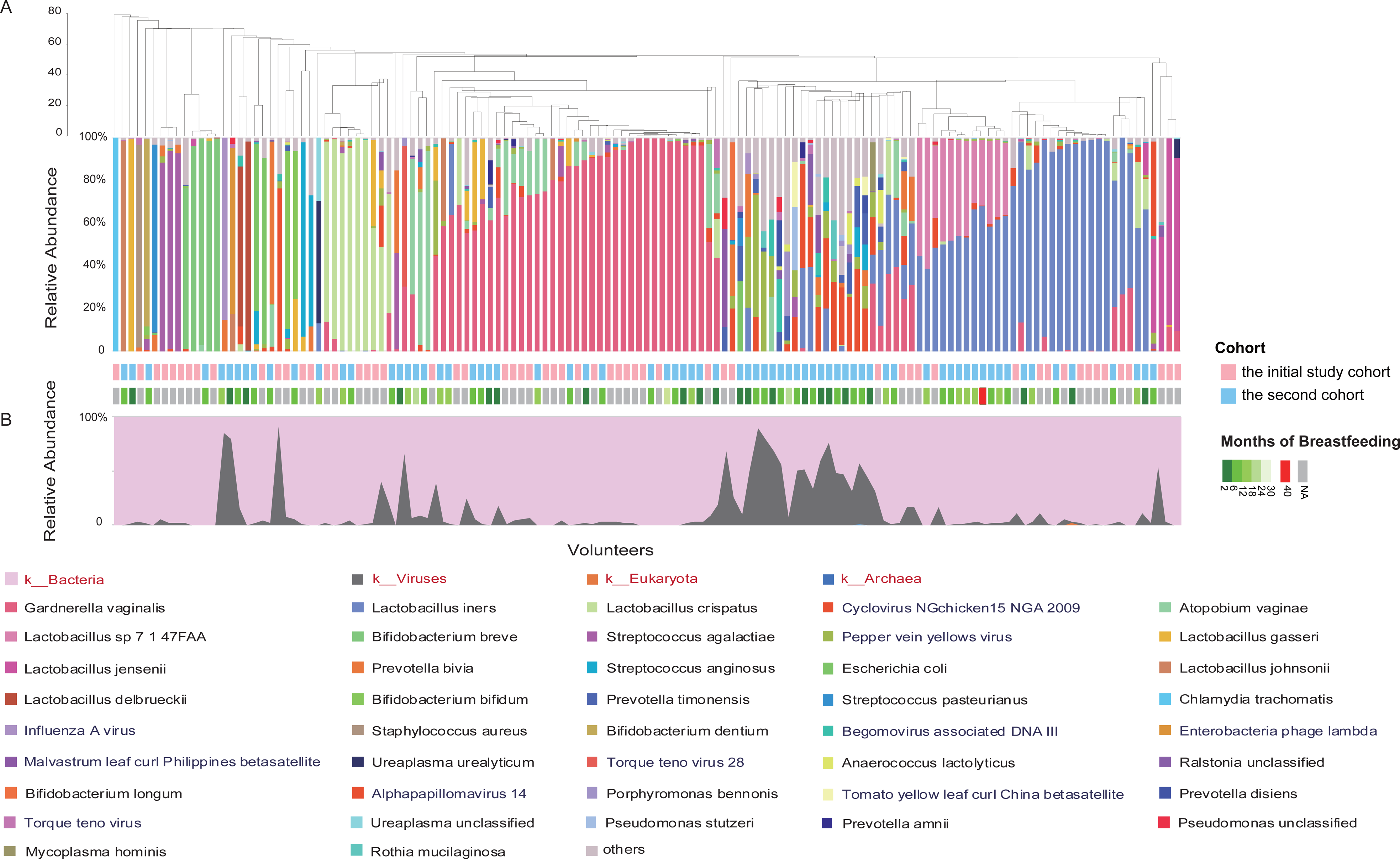
Vagino-cervical microbiome of 137 current breastfeeding women from two cohorts. The microbial composition in each sample at the species level (A) and kingdom level (B) according to MetaPhlAn2 is shown. The dendrogram in (A) was a result of a centroid linage hierarchical clustering based on Euclidean distances between the microbial composition proportion. The bottom portion of the (A) illustrates months of breastfeeding by the time of sampling for each subject and the source of each sample, and the max marked by red which is 40 months in the bar. Due to months of breastfeeding is only available in the second cohort, the grey bars mean lack of this data. The red, black and blue labels used in legend denote kingdom, bacteria, and viruses, respectively.

45 of the volunteers in the second cohort were postmenopausal (median age 54, 95% CI: 52-55), an age group untouched by HMP. Metagenomic shotgun data revealed diminished Lactobacilli in their vagino-cervical microbiome, while the mean proportion of viral sequences reached 37% (Figure S3). HPV was not the most abundant or prevalent virus in these postmenopausal individuals; a diverse range of viruses named after plant or animal hosts could be detected. In contrast, there was no sign of fungal boom. The non-Lactobacilli bacterial species were also known as vaginal or oral species, with overgrowth of *Escherichia* in only 2% of the individuals (Figure S3C).The taxonomic profile remained robust when we arbitrarily trimmed the paired-end 100bp data to single-end 50bp or single-end 100bp (Spearman’s correlation coefficient = 0.998 between Pseudo-SE50 and Pseudo-SE100, 0.993 between Pseudo-SE50 and PE100, Figure S4).

As a whole, the vagino-cervical microbial composition showed the greatest explained variances for these questionnaire data collected for the female reproductive tract samples, followed by other data collected on the same day, such as fecal microbiome composition, psychological questionnaire, plasma metabolites, immune indices, facial skin imaging and medical test data (Figure 5A). *L. crispatus* in the fecal microbiome was most predictive of the vagino-cervical microbiome, and plasma phosphoserine, L-homocitrulline, testosterone were ranked at the top among metabolites (largest Spearman’s cc between RVCV prediction and observed data, and p < 0.01 with 999 permutations, Figure 5B), while weaker signals were observed for self-assessed mental status, and pores on the cheeks (p < 0.05 but q > 0.1, Figure 5B).

**Figure 5.**
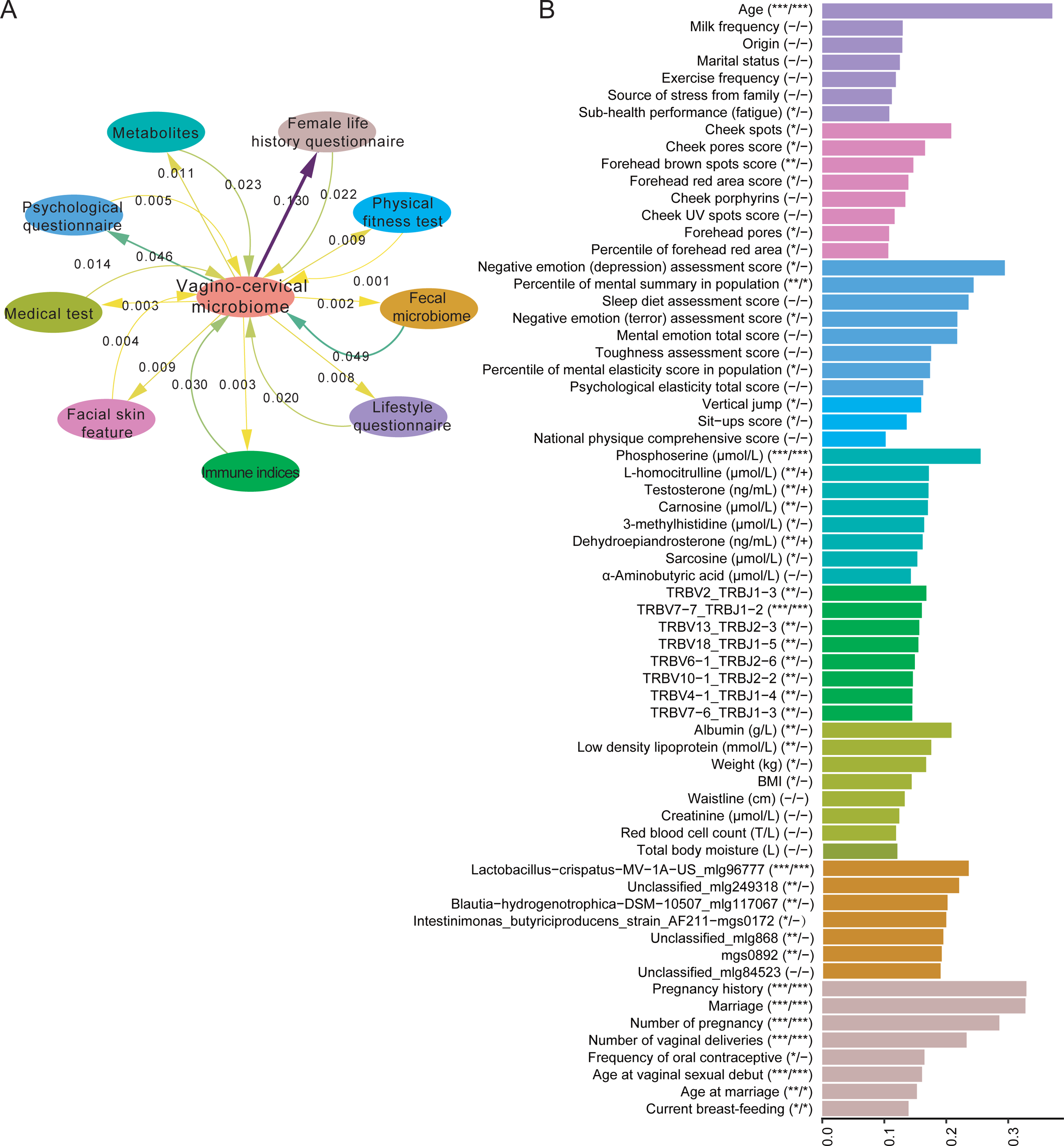
Global view of factors influencing the vagino-cervical microbiome in the initial study cohort. (A) Predicting the vagino-cervical microbiome from each omics and vice versa using stepwise redundancy analysis. Numbers on the straight arrows indicate adjusted R-squared from vagino-cervical microbiome to omics data; numbers on the curved arrows indicate adjusted R-squared from omics data to vagino-cervical microbiome. (B) Top 8 factors in each type of omics that are predicted by the vagino-cervical microbiome. The length of the bars represents RFCV Rcorrelation coefficient between the actual measurements and the values predicted by 5-fold cross-validated random forest models. First column stars after y-axis label are 999 times permutation p-value, second column stars are BH adjust p-value within each omics (4308 comparisons), an “+” denotes Q-value <0.1, an asterisk denotes Q-value <0.05, two asterisks denote Q-value <0.01, three asterisks denote Q-value <0.001.

### Specific influences from pregnancy histories, contraception and menstrual symptoms

Marital status was one of the most significant factors to influence the vagino-cervical microbiome (Figure 3, Figure S2). It showed negative correlations with relative abundances of *L. crispatus* (Spearman’s correlation, q value=1.71E-08), *Acinetobacter* species (q value=5.02E-07), *L. jensenii* (q value=0.0001), *L. vaginalis* (q value=0.000086), and *Ureaplasma parvum* (q value=0.012) and positive correlation with *Bifidobacterium breve* (q value=0.0818) (Figure 6A, Table S3). Compared to unmarried women, married women had higher concentrations of plasma 25-hydroxy vitamin D, and their plasma testosterone (responsible for sexual drive), dehydroepiandrosterone (DHEA), and creatinine had declined (Figure 6A). The age at vaginal sexual debut showed positive correlations with *Gardnerella vaginalis* and *Scardovia* species, and negative associations with *L. crispatus* and *L. iners* (Figure 6B, Table S3).

**Figure 6.**
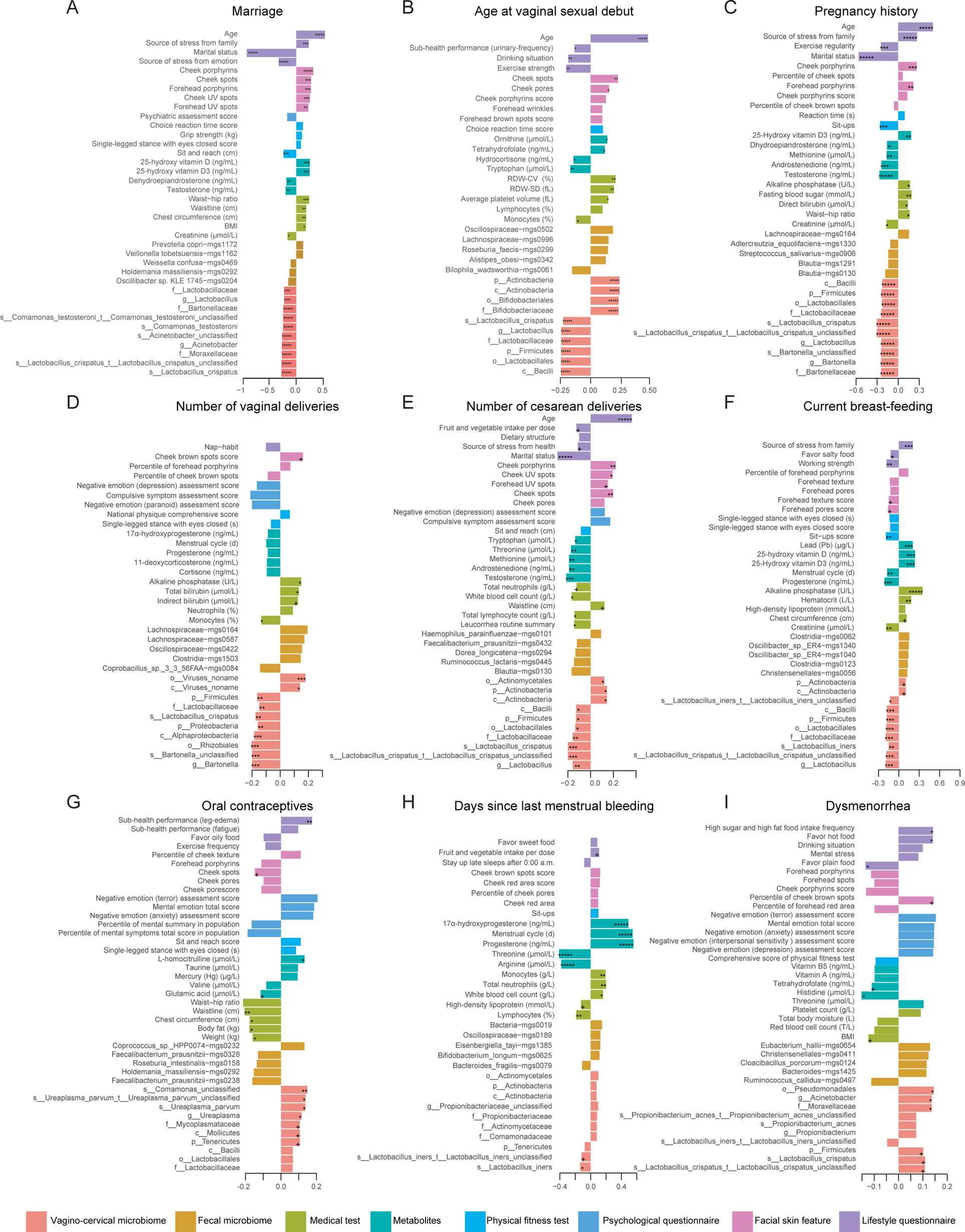
Specific influences from reproductive factors on omics data in the initial study cohort. The factors shown ((A) Marriage; (B) Age at vaginal sexual debut; (C) Pregnancy history; (D) Number of vaginal deliveries; (E) Number of cesarean deliveries; (F) Current breast-feeding; (G) Oral contraceptives; (H) Days since last menstrual bleeding; (I) Dysmenorrhea). The bars are colored according to the type of omics, as in Figure 3 The metabolites such as amino acids, hormones and vitamins are measured in plasma, and trace elements are measured in whole blood. The blood biochemistry such as alkaline phosphatase, fasting blood sugar, direct bilirubin, creatinine, total bilirubin, high-density lipoprotein is measured in serum. The length of the bars represents Spearman’s correlation coefficient between the factor and the omics data. A plus sign denotes BH-adjusted P-value <0.1 within each omics (average rank selects 13425 edges from 66301 associations, 523 edges for vagino-cervical microbiome, 11775 for fecal microbiome, 321 for metabolites, 280 for medical test, 70 for physical fitness test, 137 for facial skin feature, 319 for lifestyle questionnaire), an asterisk denotes Q-value <0.05, two asterisks denote Q-value <0.01, three asterisks denote Q-value <0.001, four asterisks denote Q-value <0.0001, and five asterisks denote Q-value <0.00001.

Similarly, the women who went through pregnancy showed relatively less *Lactobacillus* compared to nullipara, including *L. crispatus*, *L. vaginalis*, and *L. jensenii*, but were enriched for *Actinobacteria*, *Bifidobacteriaceae*, *Anelloviridae* and *Torque teno* virus (the most abundant anellovirus in the human virome) in the vagino-cervical microbiome (Figure 6C, Table S3, Figure S5). We observed increased concentrations of plasma vitamin D, blood glucose and direct bilirubin in women with previous pregnancy, but lower concentrations of plasma testosterone, androstenedione, dehydroepiandrosterone and methionine (Figure 6C). Vaginal deliveries were associated with decreased *L. crispatus*, *L. jensenii*, *Bartonella* species, *Prevotella timonensis*, and increased *Lactobacillus* sp. 7_1_47FAA (Figure 6D, Table S3). We observed a lower relative abundance of *Ureaplasma parvum* in individuals who delivered by cesarean section (Figure 6E, Table S3), a bacterium commonly isolated from pregnant women (Anderson et al., 2013) and recently reported in the lower respiratory tract of preterm infants (Pattaroni et al., 2018) and in preterm placenta (de Goffau et al., 2019). Compared with vaginal deliveries, the plasma concentrations of testosterone, androstenedione, methionine, threonine and tryptophan were significantly lower in women who experienced caesarean section (Figure 6D, 6E). In addition, these women more often suffered from an abnormal leucorrhoea (Figure 6E).

63 volunteers in our initial cohort and 74 in the second cohort happened to be actively breast-feeding. *L. crispatus* type which abundance >50% was found to be less (5.8%) in these actively breast-feeding individuals, in contrast to *G. vaginalis* type which was found in 25.5% of these individuals (Figure 6F, Figure 4). Alkaline phosphatase, plasma vitamin D and lead (Pb) were found to be more abundant in the breast-feeding individuals, consistent with slight reduction of progesterone and creatinine levels in lactating women (Figure 6F). Gravida and para, especially deliveries by caesarean section, appeared associated with lower facial skin score ranking in population such as UV spots and porphyrins, but these problems were less pronounced in caesarean section mothers with breast-feeding (Figure 6C-6F).

We found multiple significant associations between the contraceptive methods of participants and their vagino-cervical microbiome. Condom usage showed negative correlations with *L. iners* and *Comamonas* species, and positive associations with *L. gasseri* (Table S2, Figure 7, Table S3). Oral contraceptives, although still rare in our cohort, were associated with increased *Ureaplasma parvum and Comamonas* species (Table S2, Figure 6G, Table S3, and Figure 7). *Comamonas* has been implicated in fecundity in *C. elegans* and identified as a marker for infertility due to endometriosis (Chen et al., 2017; MacNeil et al., 2013). *Comamonas testosteroni* TA441 is known to degrade testosterone (Horinouchi et al., 2003). Moreover, oral contraceptives exhibited a positive correlation with plasma homocitrulline, and as yet not significant declines in mental health and butyrate-producing fecal microbiota species such as *Faecalibacterium prausnizi* and *Roseburia intestinalis* (Figure 6G), which could potentially contribute to the known side effect of oral contraceptives in cardiovascular risk. Thus, although lack of contraception could lead to infection, none of the contraceptive methods appeared healthy for the microbiome in the long run.

**Figure 7.**
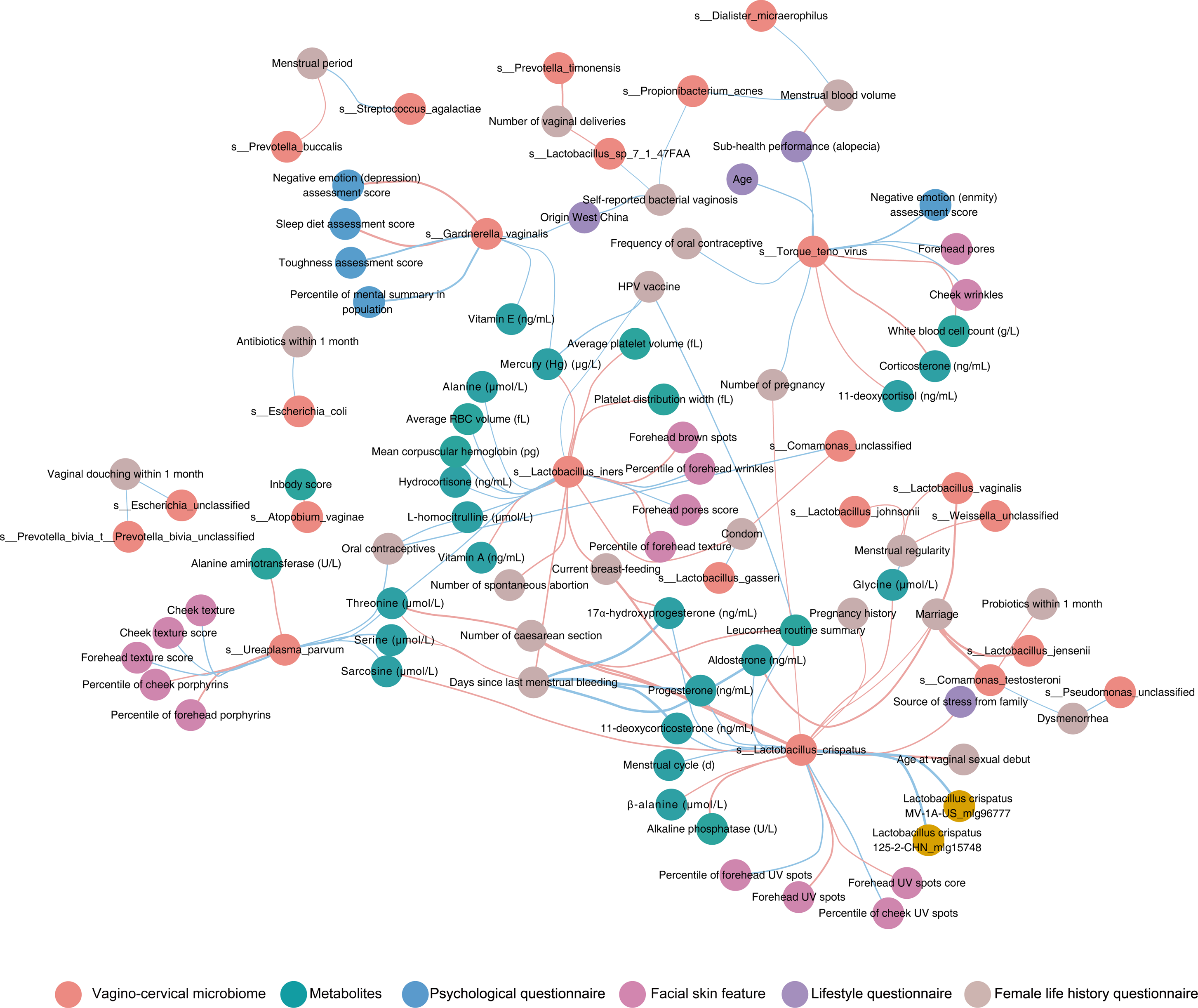
Wisdom of the crowds for the association network between vagino-cervical microbial species and other omics data in the initial study cohort. Results from generalized linear model with penalty (cv. glmnet), random forest (RFCV) and Spearman’s correlation are integrated and then visualized in CytoScape. Red lines, negative associations; cyan lines, positive associations (23385 are vagino-cervical microbiome and other omics, among them, 878 for metabolites, 785 for medical test, 216 for physical fitness test, 392 for facial skin feature, 162 for psychological questionnaire, 1096 for lifestyle questionnaire, 19856 for fecal microbiome, adding 66301 edges from female life history questionnaire and other omics, total 89828 associations).

Menstrual phases are known to influence the microbiota in the female reproductive tract (Chen et al., 2017; Gajer et al., 2012). We confirmed in our cohort the shift in *L. crispatus* and *L. iners* during menstrual cycle, coinciding with dynamics in progesterone as well as threonine and arginine (Figure 6H, Figure S6). White blood cell counts (WBC) also recovered after menses, consistent with greatest susceptibility to infection at the onset of menses. *L. vaginalis*, *L. johnsonii* and *Weissella* species showed negative correlations with self-reported regular periods (Figure 7). *Propionibacterium acnes* and *Dialister micraerophilus* were positively associated with a heavier flow (Figure 7, Table S3). Many women experience dysmenorrhea during menses (No: n=84, Slight: n=360, Serious: n=68, Table S1A). Individuals with dysmenorrhea were enriched for Pseudomonadales, *Acinetobacter* and *Moraxellaceae*, while lower in plasma level of histidine (Figure 6I), consistent with these bacteria encoding histidine decarboxylases to convert histidine into histamine (Cundell et al., 1991).

### Association between omics

Integrated association network using a wisdom of crowds approach (Marbach et al., 2012) also revealed interesting patterns (Spearman’s correlation, random forest and linear regression, with arcsin sqrt-transform for the microbiome profiles, Figure 7, Table S3). The relative abundance of *G. vaginalis* negatively associated with sleep and appetite, and mental state, while positively associated with plasma levels of mercury (Hg) and vitamin E (Figure 7, Table S3). *L. iners* negatively correlated with times of spontaneous abortion, days since last menstrual bleeding, condom usage and the plasma concentration of vitamin D and vitamin A, and positively correlated with plasma concentrations of hemoglobin, and alanine. In addition, vaccination history, such as those against HPV may be associated with higher relative abundance of *L. iners* (Figure 7, Table S3).

While lacking significant association with the vagino-cervical microbial composition, the immune repertoire data, especially TRBV7.3 and TRBV7.4, showed multiple associations with functional pathways in the vagino-cervical microbiome, such as purine and pyrimidine metabolism, synthesis of branched-chain amino acids, histidine and arginine (Figure S7). Red blood cell counts, plasma vitamin A and plasma hydroxyl vitamin D levels negatively associated with CDP-diacylglycerol biosynthesis pathways, consistent with presence of diacylglycerol kinase in Lactobacilli with anti-inflammatory functions (Andrada et al., 2017; Ganesh et al., 2017). Fecal *Coprococcus comes*, a bacterium previously reported to associate with cytokine response to *Candida albicans* (Schirmer et al., 2016), was seen here to associate with isoleucine pathways in the vagino-cervical microbiome (Figure S7).

The second cohort had saliva metagenomics data (Figure 1), which allowed us to explore the oral microbiome, a site that was reported to show similarity to the placenta microbiome (Aagaard et al., 2014). Associations with the vagino-cervical microbiome could be observed (Table S4). Oral human herpesvirus was positively correlated with Bifidobacteriaceae and *G. vaginalis* in vagina. *Scardovia wiggsiae*, a bacterium previously reported to associate with early childhood caries, showed positive correlation with vaginal *Staphylococcus* species. Oral *Treponema lecithinolyticum* showed positive correlation with *Dialister micraerophilus*.

We next analyzed the integrated association network between omics in the second cohort (Figure S8), and patterns consistent with the initial cohort could be identified (Table S4B). Pregnancy history, current breast-feeding showed negative correlations with relative abundances of *L. crispatus*, *L. jensenii*, *L. vaginalis*, *L. iners* and *U. parvum*.

We also reconfirmed a lower relative abundance of *U. parvum* and a higher abundance of *P. timonensis* in individuals who delivered by cesarean section, while individuals with past vaginal deliveries showed a lower relative abundance of *P. disiens, P. timonensis, P. buccalis* and *Finegoldia magna* (Table S4B, q<0.1). Higher concentrations of dehydroepiandrosterone, testosterone were observed in women with higher abundance of *L. crispatus*. *Finegoldia magna* negatively correlated with Vitamin B1. *Peptoniphilus harei* negatively correlated with testosterone. *P. bivia* negatively correlated with progesterone. We observed higher concentrations of threonine, arginine, and serine, in women with higher abundance of *U. parvum*, *A. vaginae* and *G. vaginalis* positively correlated with the red blood cell (RBC)*. Finegoldia magna* and *Peptoniphilus harei* positively correlated with the urine specific gravity, used in the evaluation of kidney function*. L. crispatus* show the strongest positive correlation with gut microbe. The BV species *P. bivia* negatively associated with fecal *Butyricimonas* species and *Clostridia* species, while positively associated with fecal *P. bivia*, suggesting a fecal reservoir for recurrent vaginal infections. *P. disiens, P. buccalis* and *P. timonensis* negatively associated with physical fitness test score (Figure S8). The wrinkles and red area in forehead were relieved in individuals who enriched for *L. iners*. Together, these results confirmed the long-range links between the vagino-cervical microbiome and other body sites.

## DISCUSSION

As the largest metagenomic study for the vagino-cervical microbiome, our data revealed lesser known subtypes for the vaginal microbiota, and put into context viral and fungal sequences present in healthy women. Our multi-omic data could help target efforts aimed at promoting a healthy reproductive tract microbiota and offering better advices for mothers from pregnancy to recovery, as well as preventing infections from viruses such as HIV and HPV. For example, whether vitamin D supplementation should be considered in African countries to reduce vaginal *L. iners* or rise *L. crispatus*, as was tried recently in a pregnant cohort (Jefferson et al., 2019). Gut probiotics such as *L. casei*, *B. longum* have been reported to increase vitamin D in ovariectomy-induced mice model of osteoporosis (Montazeri-Najafabady et al., 2018), and our results suggest that there might be counterparts in the female reproductive tract after getting married and pregnant.

Recent publications from iHMP (integrative Human Microbiome Project) provided longitudinal data during pregnancy and showed relatively more *L. iners* compared to non-pregnant individuals even in women of African American history (DiGiulio et al., 2015; Fettweis et al., 2019; Petricevic et al., 2014; Serrano et al., 2019). In our data, the *L. iners* versus *L. crispatus* shift was apparent in women with past pregnancy, and *L. iners* interestingly showed a negative association with number of spontaneous abortions (Figure 7). *L. iners* and other Lactobacilli have been detected in the placenta (Aagaard et al., 2014; de Goffau et al., 2019; Lannon et al., 2019; Macklaim et al., 2011; Seferovic et al., 2019), amniotic fluid, nasal and pharyngeal sites (Boeck et al.; Wang et al., 2018a). Aldosterone, a major mineralocorticoid for which we observe association with potentially beneficial bacteria in the gut microbiome in an accompanying study (Jie et al., 2019), positively associated with *L. crispatus* in the vagino-cervical microbiome, while a precursor for aldosterone, corticosterone, positively associated with *L. iners* (Figure 7). How vitamin D and hormone metabolism might impact the vagino-cervical microbiome would require further studies. How the uterus and the microbiome recover during breastfeeding is also of interest for both the mother and future children (Anton et al., 2018; DiGiulio et al., 2015). *S. anginosus*, has been shown to be more abundant in the gut microbiome of individuals with atherosclerotic cardiovascular diseases (Jie et al., 2017), and here we see young women who harbored > 50% *S. anginosus* in cervical samples showing higher plasma creatinine and lower 17α -hydroxyprogesterone, which α might relate to preterm birth (de Goffau et al., 2019). The vagino-cervical microbiome of postmenopausal women revealed a myriad of viral and bacterial species. While fungal growth might be unfavorable both due to high pH and lack of glycans. While the focus in the field has always been infection, our data highlight major aspects that are worth further investigations for women in the modern world.

While samples from multiple body sites (peritoneal fluid from the pouch of Douglas, fallopian tubes, endometrium, cervical mucus and two vaginal sites) were taken from volunteers with benign conditions such as hysteromyoma, adenomyosis and endometriosis (Chen et al., 2017; Li et al., 2018), we were only able to sample the vagino-cervical microbiome in this healthy cohort. Interestingly, our cohort generally lacked bacteria known for BV (e.g. (Fettweis et al., 2019; Serrano et al., 2019).) and also contained less *Prevotella* compared to some of the surgically sampled individuals.

In the postmenopausal samples, some of the bacteria previously reported in the upper reproductive tract (Chen et al., 2017; Lee et al., 2019; Li et al., 2018) and involved in degradation of hormones (Wang et al., 2018b), e.g. *Pseudomonas* spp. could be seen in the vagino-cervical microbiome, while disease relevance such as *A. vaginae* and *Porphyromonas* in endometrial cancer would require more evidence (Walther-António et al., 2016)..

By reaching to the cervix with a cytobrush, reflecting both the upper and the lower reproductive tract (Chen et al., 2017; Li et al., 2018), we show here that the association with the fecal microbiome was scarce overall, and the notion of reservoirs in the intestine or other sites for vaginocervical bacteria (Marrazzo et al., 2012) would need further investigation, especially in light of individual differences in the number of CD4+ T cells in mucosal sites (Gosmann et al., 2017). There were other interesting associations between different species that potentially involve immune modulation (Fig. 7). The plasma metabolites and T cell receptor types associated with the vagino-cervical microbiome are distinct from those associated with the fecal microbiome (Jie et al., 2019). Vaginal *Prevotella* could induce more CD4+ T cells (Gosmann et al., 2017), but the *Prevotella* species are not the dominant gut species of *Prevotella copri* which may compete with *Bacteroides* spp. (Jie et al., 2017). The vagino-cervical microbiome also better predicted facial skin features compared to the fecal microbiome (Jie et al., 2019), perhaps due to a clearer pattern of hormone and immune signatures. The associations with physical fitness tests and self-reported physical activities were however less prominent in the vagino-cervical compared to the fecal microbiome (Jie et al., 2019), as the changes due to pregnancy, delivery and breastfeeding may not be easily modifiable with physical activity. As a densely populated microbiota other than the distal gut, the vagino-cervical microbiota has the potential to reflect or even influence physiology elsewhere in the human body.

## ACKNOWLEDGMENTS

This research was supported by the Shenzhen Municipal Government of China (JCYJ20170817145523036) and Shenzhen Peacock Plan (No. KQTD20150330171505310). The authors are very grateful to colleagues at BGI-Shenzhen for sample collection, DNA extraction, library construction, sequencing, and discussions. We thank Dr. Xin Mu and Dr. Kou Qin for helpful comments.

## AUTHOR CONTRIBUTIONS

H.J. and C.C. conceived and organized this study. J.W. initiated the overall health project. C.C., L.H., F.L, L.S., Y.L., X.Y. and N.L. performed the sample collection and questionnaire collection. X.T., Y.K., W.R., Y.L., D.Z., X.Q., X.C., and J.Z. provided the omics data. Z.J., C.C., F.L, L.H., and L.T. performed the bioinformatic analyses, H.J., C.C., Z.J. and X.Z. wrote the manuscript. All authors contributed to data and texts in this manuscript.

## STAR METHODS

### KEY RESOURCES TABLE

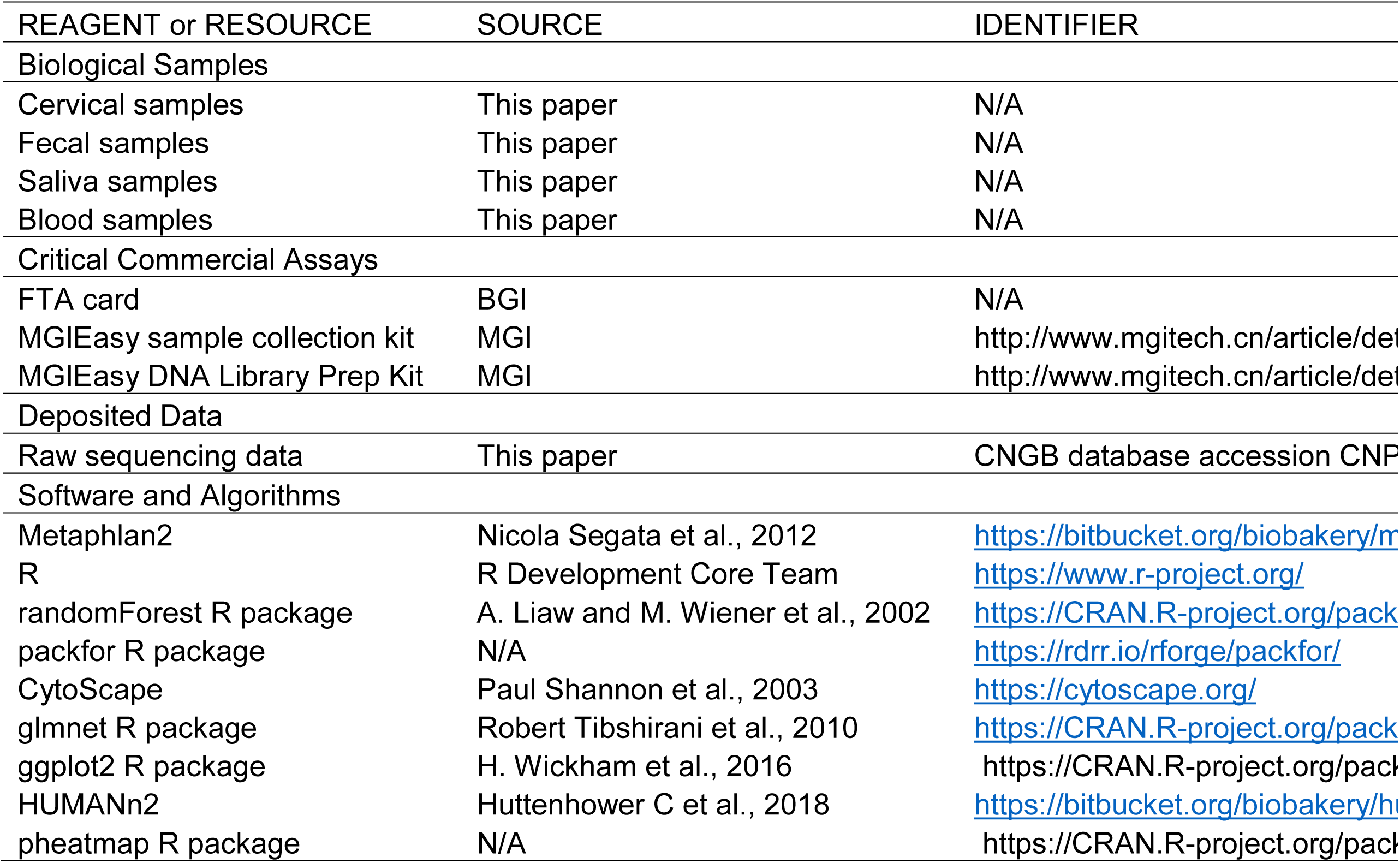

### CONTACT FOR REAGENT AND RESOURCE SHARING

Further information and requests for resources and reagents should be directed to, and will fulfilled by, the Lead Contact, Huijue Jia.

### EXPERIMENTAL MODEL AND SUBJECT DETAILS

#### Initial Study Cohort

As the first time point for the vagino-cervical microbiome of the 4D-SZ cohort, 516 volunteers joined between May 2017 and July 2017 during an annual physical examination. Baseline characteristics of the cohort are shown in Table S1A.

The study was approved by the Institutional Review Boards (IRB) at BGI-Shenzhen, and all participants provided written informed consent at enrolment.

#### The Second Cohort

An independent cohort of 632 were also recruited between May 2018 and July 2018. 2018 data for 4D-SZ volunteers who were already included in the initial cohort would be published in a future study. The process of omics data collection was consistent with the initial cohort. In addition, saliva samples were collected only in this cohort. Baseline characteristics of the cohort are shown in Table S1E.

The study was approved by the Institutional Review Boards (IRB) at BGI-Shenzhen, and all participants provided written informed consent before any material was taken.

### METHODS DETAILS

#### Demographic Data Collection

During physical examination, the volunteers received three kinds of online questionnaire. The female life history questionnaire contained pregnancy and delivery history, menstrual phases, sexual activity, contraceptive methods. The lifestyle questionnaire contained age, disease history, eating and exercise habits. The psychological questionnaire contained the evaluation of irritability, dizziness, frustration, fear, appetite, self-confidence, resilience (Table S1A, S1E).

#### Samples Collection

Cervical samples were collected and smeared in the Flinders Technology Associates (FTA) cards by the doctor during gynecological examination. Fecal samples and saliva samples were self-collected by volunteers. Cervical samples, fecal samples and saliva samples were stored for metagenomic shotgun sequencing. The overnight fasting blood samples were drawn from a cubital vein of volunteers by the doctors.

#### DNA extraction and metagenomic shotgun sequencing

DNA extraction of cervical samples and fecal samples was performed as described (Chen et al., 2017; Li et al., 2014). Metagenomic sequencing was done on the BGISEQ-500 platform, which is highly comparable to Illumina HiSeq platforms in metagenomic and other sequencing applications (Fang et al., 2018; Han et al., 2018; Li et al., 2018; Pan et al., 2018). 50bp of singled-end reads for cervical samples collected in the initial study cohort, and on average 208.76 million raw reads were sequenced for each sample (Table S1B); 100bp of singled-end reads for fecal samples collected in the initial study cohort, and on average 85.63 million raw reads were sequenced for each sample (Table S1B); 100bp of paired-end reads for cervical samples, fecal samples and saliva sample collected in the second cohort, and on average 158.91 million raw reads, 75.84 million raw reads were sequenced for each cervical sample and saliva sample, respectively (Table S1B). Quality control and alignment to GRCh38 was performed as previously described (Fang et al., 2018; Li et al., 2018).

#### UHPLC-MS quantification of amino acids

40 µl plasma was deproteinized with 20 µl 10% (w/v) sulfosalicylic acid (Sigma) containing internal standards, then 120 µl aqueous solution was added. After centrifuged, the supernatant was used for analysis. The analysis was performed by ultra high pressure liquid chromatography (UHPLC) coupled to an AB Sciex Qtrap 5500 mass spectrometry (AB Sciex, US) with the electrospray ionization (ESI) source in positive ion mode. A Waters ACQUITY UPLC HSS T3 column (1.8 µm, 2.1 × 100 mm) was used for amino compound separation with a flow rate at 0.5 ml/min and column temperature of 55 °C. The mobile phases were (A) water containing 0.05% and 0.1% formic acid (v/v), (B) acetonitrile containing 0.05% and 0.1% formic acid (v/v). The gradient elution was 2% B kept for 0.5 min, then changed linearly to 10% B during 1 min, continued up to 35% B in 2 min, increased to 95% B in 0.1 min and maintained for 1.4 min. Multiple Reaction Monitoring (MRM) was used to monitor all amino compounds. The mass parameters were as follows, Curtain gas flow 35 L/min, Collision Gas (CAD) was medium, Ion Source Gas 1 (GS 1) flow 60 l/min, Ion Source Gas 2 (GS2) flow 60 l/min, IonSpray Voltage (IS) 5500V, temperature 600 °C. All amino compound standards were purchased from sigma and Toronto research chemical (TRC).

#### UHPLC-MS quantification of Hormones

250 µl plasma was diluted with 205 µl aqueous solution, For SPE experiments, HLB (Waters, USA) was activated with 1.0 ml of dichloromethane, acetonitrile, methanol, respectively and was equilibrated with 1.0 ml of water. The pretreated plasma sample was loaded onto the cartridge and was extracted using gravity. Clean up was accomplished by washing the cartridges with 1.0 ml of 25% methanol in water. After drying under vacuum, samples on the cartridges were eluted with 1.0 ml of dichloromethane. The eluted extract was dried under nitrogen and the residual was dissolved with 25% methanol in water and was transferred to an autosampler vial prior to LC–MS/MS analysis. The analysis was performed by UHPLC coupled to an AB Sciex Qtrap 5500 mass spectrometry (AB Sciex, US) with the atmospheric pressure chemical ionization (APCI) source in positive ion mode. A Phenomone Kinetex C18 column (2.6 µm, 2.1 × 50 mm) was used for steroid hormone separation with a flow rate at 0.8 ml/min and column temperature of 55 °C. The mobile phases were (A) water containing 1mM Ammonium acetate, (B) Methanol containing 1mM Ammonium acetate. The gradient elution was 25% B kept for 0.9min, then changed linearly to 40% B during 0.9min, continued up to 70% B in 2 min, increased to 95% B in 0.1 min and maintained for 1.6 min. Multiple Reaction Monitoring (MRM) was used to monitor all steroid hormone compounds. The mass parameters were as follows, Curtain gas flow 35 l/min, Collision Gas (CAD) was medium, Ion Source Gas 1 (GS 1) flow 60 l/min, Ion Source Gas 2 (GS 2) flow 60 l/min, Nebulizer Current (NC) 5, temperature 500 °C. All steroid hormone profiling compound standards were purchased from sigma, Toronto research chemical (TRC), Cerilliant and DR. Ehrenstorfer.

#### ICP-MS quantification of trace elements

200 µl of whole blood were transferred into a 15 mL polyethylene tube and diluted 1:25 with a diluent solution consisting of 0.1% (v/v) Triton X-100, 0.1% (v/v) HNO_3_,2mg/L AU, and internal standards (20 µg/L). The mixture was sonicated for 10min before ICP-MS analysis. Multi-element determination was performed on an Agilent 7700x ICP-MS (Agilent Technologies, Tokyo, Japan) equipped with an octupole reaction system (ORS) collision/reaction cell technology to minimize spectral interferences. The continuous sample introduction system consisted of an autosampler, a quartz torch with a 2.5-mmdiameter injector with a Shield Torch system, a Scott double-pass spray chamber and nickel cones (Agilent Technologies, Tokyo, Japan). A glass concentric MicroMist nebuliser (Agilent Technologies, Tokyo, Japan) was used for the analysis of diluted samples.

#### UPLC-MS quantification of water-soluble vitamins

200 µl plasma were deproteinized with 600 µl methanol (Merck), water, acetic acid (9:1:0.1) containing internal standards, thiamine-(4-methyl-13C-thiazol-5-yl-13C3) hydrochloride (Sigma-Aldrich), levomefolic acid-13C, d3, riboflavin-13C,15N2, 4-pyridoxic acid-d3 and pantothenic acid-13C3,15N hemi calcium salt (Toronto Research Chemicals). 500 µl supernatant were dried by nitrogen flow. 60 µl water were added to the residues, vortexed, the mixture was centrifuged and the supernatant was for analysis. The analysis was performed by UPLC coupled to a Waters Xevo TQ-S Triple Quad mass spectrometry (Waters, USA) with the electrospray ionization (ESI) source in positive ion mode. A Waters ACQUITY UPLC HSS T3 column (1.7 µm, 2.1 × 50 mm) was used for water-soluble vitamins separation with a flow rate at 0.45 ml/min and column temperature of 45 °C. The mobile phases were (A) 0.1 % formic acid in water, 0.1% formic acid in methanol. The following elution gradient was used: 0–1 min,99.0%–99.0% A; 1–1.5 min, 99.0% A–97.0% A; 1.5–2 min, 97.0% A–70.0% A,2– 3.5 min, 70% A–30% A; 3.5–4.0 min, 30%A–10.0%A; 4.0–4.8 min, 10%A–10.0%A; 4.9–6.0 min, 99.0%A–99.0%A. Multiple Reaction Monitoring (MRM) was used to monitor all water-soluble vitamins. The mass parameters were as follows, the capillary voltages of 3000V and source temperature of 150°C were adopted. The desolvation temperature was 500°C. The collision gas flow was set at 0.10 ml/min. The cone gas and desolvation gas flow were 150 l/h and 1000 l/h, respectively. All water-soluble vitamins standards were purchased from Sigma-Aldrich (USA).

#### UPLC-MS quantification of fat-soluble vitamins

250 µl plasma were deproteinized with 1000 µl methanol and acetonitrile, (v/v,1:1) (Fisher Chemical) containing internal standards, all-trans-Retinol-d5, 25-HydroxyVitamin-D2-d6, 25-HydroxyVitamin-D3-d6, vitamin K1-d7, α-Tocopherol-d6 (Toronto Research Chemicals). 900 µl supernatant were dried by nitrogen flow. 80 µl 80% acetonitrile were added to the residues, vortexed, the mixture was centrifuged, and the supernatant was used for analysis. The analysis was performed by UPLC coupled to an AB Sciex Qtrap 4500 mass spectrometry (AB Sciex, USA) with the atmospheric pressure chemical ionization (APCI) source in positive ion mode. A Waters ACQUITY UPLC BEH C18 column (1.7 µm, 2.1 × 50 mm) was used for fat-soluble vitamins separation with a flow rate at 0.50 ml/min and column temperature of 45 °C. The mobile phases were (A) 0.1 % formic acid in water, (B) 0.1% formic acid in acetonitrile. The following elution gradient was used: 0–0.5 min,60.0%–60.0% A; 0.5–1.5 min, 60.0% A– 20.0% A; 1.5–2.5 min, 20.0% A–0% A,2.5–4.5 min, 0% A–0% A; 4.5–4.6 min, 0%A–60.0%A; 4.6–5.0 min, 60.0%A–60.0%A. Multiple Reaction Monitoring (MRM) was used to monitor all fat-soluble vitamins. The mass parameters were as follows, Curtain gas flow 30 l/min, Collision Gas (CAD) was medium, Ion Source Gas 1 (GS 1) flow 40 l/min, Ion Source Gas 2 (GS 2) flow 50 l/min, Nebulizer Current (NC) 5, temperature 400 °C. All fat-soluble vitamins standards were purchased from Sigma-Aldrich (USA), Toronto research chemical (TRC).

#### Sequencing of the TCR CDR3 immune repertoire

10 ml whole blood was centrifuged at 3,000 r/min for 10 min, then 165 µl buffy coat were obtained to extract DNA using MagPure Buffy Coat DNA Midi KF Kit (Magen, China). The DNA was sequenced on the BGISEQ-500 platform using 200 bp singled-end reads. The data processing was performed using Immune IMonitor (Zhang et al., 2015a).

#### Medical Parameters

All the volunteers were recruited during the physical examination. The medical test including blood tests, urinalysis, routine examination of cervical secretion. All the medical parameters were measured by the physical examination center and shown in Table S1A and Table S1E.

#### Facial skin features

The volunteers were required to clean their face without makeup after they got up in the morning. The volunteer’s frontal face was photographed by VISIA-CRTM imaging system (Canfield Scientific, Fairfield, NJ, USA) equipped with chin supports and forehead clamps that fix the face during the photographing process and maintain a fixed distance between the volunteers and the camera at all times. Eight indices were obtained including spots, pores, wrinkles, texture, UV spots, porphyrins, brown spots and red area from the cheek and forehead, respectively (Table S1A, S1E). The percentile of index was calculated based on the index value ranked in the age-matched database (Table S1A, S1E). The higher the percentile of index, the better the facial skin appears.

#### Physical fitness test

8 kinds of tests were performed to evaluate volunteers’ physical fitness condition (Table S1A, S1E). Vital capacity was measured by HK6800-FH (Hengkangjiaye, China). Single-legged stance with eyes closed was measured by HK6800-ZL. Choice reaction time was measured by HK6800-FY. Grip strength was measured by HK6800-WL. Sit and reach was measured by HK6800-TQ. Sit-ups was measured by HK6800-YW. Step index was measured by HK6800-TJ. Vertical jump was measured by HK6800-ZT. We got a measure value from each test. Then each measure value score was assigned grades A through E based on its corresponding age-matched database.

### QUANTIFICATION AND STATISTICAL ANALYSIS

#### Quality control, Taxonomic annotation and abundance calculation

The sequencing reads of fecal samples and saliva samples were quality-controlled and then aligned to hg19 to remove human reads as described previously(Fang et al., 2018). Stringent condition for removal of host sequences was used for cervical samples(Li et al., 2018), through alignment to the GRCh38 reference.

Taxonomic assignment of the high-quality cervical metagenomic data and saliva metagenomic data were performed using MetaPhlAn2(Segata et al., 2012).

Taxonomic assignment of the high-quality fecal metagenomic data was performed using the reference gene catalog comprising 9,879,896 genes(Li et al., 2014). Taxonomy of the fecal MLGs/MGSs were then determined from their constituent genes, as previously described(Moll1 et al., 2019; Nielsen et al., 2014; Qin et al., 2012; Wang and Jia, 2016).

#### Random-forest on the influence of female life history factors

The factors in female life history questionnaire were fitted against the relative abundances of metaphlan2 profile (found in at least 10% of the samples) in the cervical samples using default parameters in the RFCV regression function from randomForest package in R. Female life history factors are dummy variables such as pregnancy history (yes, no) or frequency variables such as number of caesarean section (0, 1, 2), except age is continue variables, see table S1A. In addition to comparing the predict power across factors, we use regression model instead of classification model here.

Spearman’s correlation between measured value and 5-fold cross-validation predicted value was calculated as model performance metric, then rank the key predictable factors. P-value was done using 999 permutation test.

#### The global effect size between vagino-cervical microbiome and omics data

To evaluate the combined effect size of vagino-cervical microbiome on omics data, we used forward stepwise redundancy analysis of omics data lists on the relative abundances of metaphlan2 profile in forward.sel function in the packfor package in R. This analysis provided a global versus global association between any two omics datasets that maximize the associations by use the most predict power non-redundant predictors.

#### The factors in each type of omics predicted by vagino-cervical microbiome

The factors in each type of omics were fitted against the relative abundances of metaphlan2 profile (found in at least 10% of the samples) in the cervical samples using default parameters in the RFCV regression function from randomForest package in R. Omics data are mix of dummy variables and continue variables, see table S1A. In addition to comparing the predict power across factors, we use regression model instead of classification model here. Spearman’s correlation between measured value and 5-fold cross-validation predicted value was calculated as model performance metric, then rank the top 8 predictable factors in each type. P-value was done using 999 permutation test,

#### Transformation of metagenomics profile for composition data analysis

Vaginal cervical relative abundance profile was composition data, so we adopt arcsin sqrt-transformation to make it resemble continuous for the downstream analysis (implemented in MaAsLin software, Morgan et al 2012).

#### Wisdom of crowds for robust network construction between vagino-cervical microbial species and other omics data

A new multi-omics analyses method (Marbach et al., 2012) was used to integrated coefficient of linear regression, variance importance from randomForest and

Spearman’s correlation to construct omics flux networks and then visualized in CytoScape. The details are as indicated:

Step 1: Data processing. All categorical variables in other omics data were converted into continuous variables, and nominal variables were converted into dummy variables. Missing values were filled with median, the samples which contained more than 70% missing variables were removed. The microbial species less than10% in all the samples were also removed. Removed near zero variable variables. For linear models, variables were normalized. Outliers were defined as outside of the 95% quartiles and removed.

Step 2: Method implementation. Random forest variable importance was used to identify the most important predictor variables(Louppe et al., 2013). RFCV regression function from randomForest package in R with default parameter was used to get the 5-fold average variable importance. We calculated the Spearman’s correlation with the cor.test function in base R software. For linear regression, we considered penalty regression to overcome the sparse and co-linear problem, cv.glmnet function from glmnet package in R was first used to figure out the best lambda parameter, bootstrapping glmnet with 0.632 re-sampling was performed 100 times, then we obtained the best lambda.

Step 3: Construction of robust networks. We kept first 5 average ranks for each target variable and retained edges with Spearman’s correlation Q-value <0.1. Then ggplot package in R was used to make barplot for some representative female life history factors (Figure 6). CytoScape was also used to visualize the omics network (Figure 7). The second cohort was analyzed using the same statistical method. Combining P-value is computed using Edgington method from metap package in R. Benjamini and Hochberg methods were used to adjust the multiple test P-value. The correlation is identified based on BH-adjusted P-value <0.1 when the similar microbial distribution pattern shown in the initial study cohort and the second cohort.

#### Association between microbiome pathways and other omics

Pathway profile was calculated from the vagino-cervical metagenomic data using humann2. Spearman’s correlation was calculated between the relative abundance of each pathway and other numerical data collected. R package heatmap were used for visualization. Q-value <0.1 was considered as significant.

### DATA AND SOFTWARE AVAILABILITY

Metagenomic sequencing data for all samples have been deposited to the (CNGB) database under the accession code CNP0000287.

## REFERENCES

1. Aagaard, K., Ma, J., Antony, K.M., Ganu, R., Petrosino, J., and Versalovic, J. (2014). The placenta harbors a unique microbiome. Sci. Transl. Med. 6, 237ra65.

2. Anderson, B.L., Mendez-Figueroa, H., Dahlke, J.D., Raker, C., Hillier, S.L., and Cu-Uvin, S. (2013). Pregnancy-induced changes in immune protection of the genital tract: defining normal. Am. J. Obstet. Gynecol. 208, 321.e1–9.

3. Andrada, E., Liébana, R., and Merida, I. (2017). Diacylglycerol Kinase ζ Limits Cytokine-dependent Expansion of CD8+ T Cells with Broad Antitumor Capacity. EBioMedicine 19, 39–48.

4. Anton, L., Sierra, L.-J., DeVine, A., Barila, G., Heiser, L., Brown, A.G., and Elovitz, M.A. (2018). Common Cervicovaginal Microbial Supernatants Alter Cervical Epithelial Function: Mechanisms by Which Lactobacillus crispatus Contributes to Cervical Health. Front. Microbiol. 9.

5. Boeck, I. De, Broek, M.F.L. van den, Llonsius, C.N., Martens, K., Wuyts, S., and Wittouck, S. Lactobacilli Have a Niche in the Human Nose. Cell Rep.

6. Bradford, L.L., and Ravel, J. (2017). The vaginal mycobiome: A contemporary perspective on fungi in women’s health and diseases. Virulence 8, 342–351.

7. Byrd, A.L., Belkaid, Y., and Segre, J.A. (2018). The human skin microbiome. Nat. Rev. Microbiol. 16, 143–155.

8. Chen, C., Song, X., Wei, W., Zhong, H., Dai, J., Lan, Z., Li, F., Yu, X., Feng, Q., Wang, Z., et al. (2017). The microbiota continuum along the female reproductive tract and its relation to uterine-related diseases. Nat. Commun. 8, 875.

9. Cundell, D.R., Devalia, J.L., Wilks, M., Tabaqchali, S., and Davies, R.J. (1991). Histidine decarboxylases from bacteria that colonise the human respiratory tract. J. Med. Microbiol. 35, 363–366.

10. DiGiulio, D.B., Callahan, B.J., McMurdie, P.J., Costello, E.K., Lyell, D.J., Robaczewska, A., Sun, C.L., Goltsman, D.S.A., Wong, R.J., Shaw, G., et al. (2015). Temporal and spatial variation of the human microbiota during pregnancy. Proc. Natl. Acad. Sci. 112, 11060–11065.

11. Fang, C., Zhong, H., Lin, Y., Chen, B., Han, M., Ren, H., Lu, H., Luber, J.M., Xia, M., Li, W., et al. (2018). Assessment of the cPAS-based BGISEQ-500 platform for metagenomic sequencing. Gigascience 7, 1–8.

12. Fettweis, J.M., Serrano, M.G., Brooks, J.P., Edwards, D.J., Girerd, P.H., Parikh, H.I., Huang, B., Arodz, T.J., Edupuganti, L., Glascock, A.L., et al. (2019). The vaginal microbiome and preterm birth. Nat. Med. 25, 1012–1021.

13. Fredricks, D.N., Fiedler, T.L., and Marrazzo, J.M. (2005). Molecular Identification of Bacteria Associated with Bacterial Vaginosis. N. Engl. J. Med. 353, 1899–1911.

14. Gajer, P., Brotman, R.M., Bai, G., Sakamoto, J., Schutte, U.M.E., Zhong, X., Koenig, S.S.K., Fu, L., Ma, Z., Zhou, X., et al. (2012). Temporal Dynamics of the Human Vaginal Microbiota. Sci. Transl. Med. 4, 132ra52–132ra52.

15. Ganesh, B.P., Hall, A., Ayyaswamy, S., Nelson, J.W., Fultz, R., Major, A., Haag, A., Esparza, M., Lugo, M., Venable, S., et al. (2017). Diacylglycerol kinase synthesized by commensal Lactobacillus reuteri diminishes protein kinase C phosphorylation and histamine-mediated signaling in the mammalian intestinal epithelium. Mucosal Immunol.

16. de Goffau, M.C., Lager, S., Sovio, U., Gaccioli, F., Cook, E., Peacock, S.J., Parkhill, J., Charnock-Jones, D.S., and Smith, G.C.S. (2019). Human placenta has no microbiome but can contain potential pathogens. Nature.

17. Gosmann, C., Anahtar, M.N., Handley, S.A., Farcasanu, M., Abu-Ali, G., Bowman, B.A., Padavattan, N., Desai, C., Droit, L., Moodley, A., et al. (2017). Lactobacillus-Deficient Cervicovaginal Bacterial Communities Are Associated with Increased HIV Acquisition in Young South African Women. Immunity 46, 29–37.

18. Han, M.M., Hao, L., Lin, Y., Li, F., Wang, J., Yang, H., Xiao, L., Kristiansen, K., Jia, H., and Li, J. (2018). A novel affordable reagent for room temperature storage and transport of fecal samples for metagenomic analyses. Microbiome 6, 43.

19. Horinouchi, M., Hayashi, T., Yamamoto, T., and Kudo, T. (2003). A new bacterial steroid degradation gene cluster in Comamonas testosteroni TA441 which consists of aromatic-compound degradation genes for seco-steroids and 3-ketosteroid dehydrogenase genes. Appl. Environ. Microbiol. 69, 4421–4430.

20. Jefferson, K.K., Parikh, H.I., Garcia, E.M., Edwards, D.J., Serrano, M.G., Hewison, M., Shary, J.R., Powell, A.M., Hollis, B.W., Fettweis, J.M., et al. (2019). Relationship between vitamin D status and the vaginal microbiome during pregnancy. J. Perinatol. 39, 824–836.

21. Jie, Z., Xia, H., Zhong, S.-L., Feng, Q., Li, S., Liang, S., Zhong, H., Liu, Z., Gao, Y., Zhao, H., et al. (2017). The gut microbiome in atherosclerotic cardiovascular disease. Nat. Commun. 8, 845.

22. Jie, Z., Liang, S., Ding, Q., Tang, S., Wang, D., Zhong, H., Jia, H., and Xu, X. (2019). A multi-omic cohort as a reference point for promoting a healthy gut microbiome. To Be Submitt. Soon.

23. Lannon, S.M.R., Adams Waldorf, K.M., Fiedler, T., Kapur, R.P., Agnew, K., Rajagopal, L., Gravett, M.G., and Fredricks, D.N. (2019). Parallel detection of lactobacillus and bacterial vaginosis-associated bacterial DNA in the chorioamnion and vagina of pregnant women at term. J. Matern. Neonatal Med. 32, 2702–2710.

24. Lee, S., La, T.-M., Lee, H.-J., Choi, I.-S., Song, C.-S., Park, S.-Y., Lee, J.-B., and Lee, S.-W. (2019). Characterization of microbial communities in the chicken oviduct and the origin of chicken embryo gut microbiota. Sci. Rep. 9, 6838.

25. Li, F., Chen, C., Wei, W., Wang, Z., Dai, J., Hao, L., Song, L., Zhang, X., Zeng, L., Du, H., et al. (2018). The metagenome of the female upper reproductive tract. Gigascience 7.

26. Li, J., Jia, H., Cai, X., Zhong, H., Feng, Q., Sunagawa, S., Arumugam, M., Kultima, J.R.J.R., Prifti, E., Nielsen, T., et al. (2014). An integrated catalog of reference genes in the human gut microbiome. Nat. Biotechnol. 32, 834–841.

27. Lloyd-Price, J., Mahurkar, A., Rahnavard, G., Crabtree, J., Orvis, J., Hall, A.B., Brady, A., Creasy, H.H., McCracken, C., Giglio, M.G., et al. (2017). Strains, functions and dynamics in the expanded Human Microbiome Project. Nature.

28. Louppe, G., Wehenkel, L., Sutera, A., and Geurts, P. (2013). Understanding variable importances in forests of randomized trees. In Advances in Neural Information Processing Systems 26, C.J.C. Burges, L. Bottou, M. Welling, Z. Ghahramani, and K.Q. Weinberger, eds. (Curran Associates, Inc.), pp. 431–439.

29. Ma, B., Forney, L.J., and Ravel, J. (2012). Vaginal microbiome: rethinking health and disease. Annu. Rev. Microbiol. 66, 371–389.

30. Macklaim, J.M., Gloor, G.B., Anukam, K.C., Cribby, S., and Reid, G. (2011). At the crossroads of vaginal health and disease, the genome sequence of Lactobacillus iners AB-1. Proc. Natl. Acad. Sci. U. S. A. 108 Suppl, 4688–4695.

31. MacNeil, L.T., Watson, E., Arda, H.E., Zhu, L.J., and Walhout, A.J.M. (2013). Diet-induced developmental acceleration independent of TOR and insulin in C. elegans. Cell 153, 240–252.

32. Marbach, D., Costello, J.C., Küffner, R., Vega, N.M., Prill, R.J., Camacho, D.M., Allison, K.R., DREAM5 Consortium, Kellis, M., Collins, J.J., et al. (2012). Wisdom of crowds for robust gene network inference. Nat. Methods 9, 796–804.

33. Marrazzo, J.M., Fiedler, T.L., Srinivasan, S., Thomas, K.K., Liu, C., Ko, D., Xie, H., Saracino, M., and Fredricks, D.N. (2012). Extravaginal reservoirs of vaginal bacteria as risk factors for incident bacterial vaginosis. J. Infect. Dis. 205, 1580–1588.

34. Methé, B.A., Nelson, K.E., Pop, M., Creasy, H.H., Giglio, M.G., Huttenhower, C., Gevers, D., Petrosino, J.F., Abubucker, S., Badger, J.H., et al. (2012). A framework for human microbiome research. Nature 486, 215–221.

35. Moll1, J.M., Rausch, P., Eriksen, C., Nielsen, H.B., Feng, Q., Xia, H., and Brix, S. (2019). Pathway-based approach to classify unannotated gut bacteria associated with insulin sensitivity. Under Revis. Nat Microbiol.

36. Montazeri-Najafabady, N., Ghasemi, Y., Dabbaghmanesh, M.H., Talezadeh, P., Koohpeyma, F., and Gholami, A. (2018). Supportive Role of Probiotic Strains in Protecting Rats from Ovariectomy-Induced Cortical Bone Loss. Probiotics Antimicrob. Proteins.

37. Nielsen, H.B., Almeida, M., Juncker, A.S., Rasmussen, S., Li, J., Sunagawa, S., Plichta, D.R., Gautier, L., Pedersen, A.G., Le Chatelier, E., et al. (2014). Identification and assembly of genomes and genetic elements in complex metagenomic samples without using reference genomes. Nat. Biotechnol. 32, 822–828.

38. O’Toole, P.W., Marchesi, J.R., Hill, C., Na, Y.C., and Kim, H.S. (2017). Next-generation probiotics: the spectrum from probiotics to live biotherapeutics. Nat. Microbiol. 2, 17057.

39. Pan, H., Guo, R., Zhu, J., Wang, Q., Ju, Y., Xie, Y., Zheng, Y., Wang, Z., Li, T., Liu, Z., et al. (2018). A gene catalogue of the Sprague-Dawley rat gut metagenome. Gigascience 7.

40. Pattaroni, C., Watzenboeck, M.L., Schneidegger, S., Kieser, S., Wong, N.C., Bernasconi, E., Pernot, J., Mercier, L., Knapp, S., Nicod, L.P., et al. (2018). Early-Life Formation of the Microbial and Immunological Environment of the Human Airways. Cell Host Microbe 24, 857–865.e4.

41. Petricevic, L., Domig, K.J., Nierscher, F.J., Sandhofer, M.J., Fidesser, M., Krondorfer, I., Husslein, P., Kneifel, W., and Kiss, H. (2014). Characterisation of the vaginal Lactobacillus microbiota associated with preterm delivery. Sci. Rep. 4, 5136.

42. Qin, J., Li, Y., Cai, Z., Li, S., Zhu, J., Zhang, F., Liang, S., Zhang, W., Guan, Y., Shen, D., et al. (2012). A metagenome-wide association study of gut microbiota in type 2 diabetes. Nature 490, 55–60.

43. Ravel, J., Gajer, P., Abdo, Z., Schneider, G.M., Koenig, S.S.K., Mcculle, S.L., Ault, K., Peralta, L., and Forney, L.J. (2010). Vaginal microbiome of reproductive-age women. Proc. Natl. Acad. Sci. 108, 4680–4687.

44. Schirmer, M., Smeekens, S.P., Vlamakis, H., Jaeger, M., Oosting, M., Franzosa, E.A., Jansen, T., Jacobs, L., Bonder, M.J., Kurilshikov, A., et al. (2016). Linking the Human Gut Microbiome to Inflammatory Cytokine Production Capacity. Cell 167, 1125–1136.e8.

45. Seferovic, M.D., Pace, R.M., Carroll, M., Belfort, B., Major, A.M., Chu, D.M., Racusin, D.A., Castro, E.C.C., Muldrew, K.L., Versalovic, J., et al. (2019). Visualization of microbes by 16S in situ hybridization in term and preterm placentas without intraamniotic infection. Am. J. Obstet. Gynecol. 221, 146.e1–146.e23.

46. Segata, N., Waldron, L., Ballarini, A., Narasimhan, V., Jousson, O., and Huttenhower, C. (2012). Metagenomic microbial community profiling using unique clade-specific marker genes. Nat. Methods 9, 811–814.

47. Sender, R., Fuchs, S., and Milo, R. (2016). Are We Really Vastly Outnumbered? Revisiting the Ratio of Bacterial to Host Cells in Humans. Cell 164, 337–340.

48. Serrano, M.G., Parikh, H.I., Brooks, J.P., Edwards, D.J., Arodz, T.J., Edupuganti, L., Huang, B., Girerd, P.H., Bokhari, Y.A., Bradley, S.P., et al. (2019). Racioethnic diversity in the dynamics of the vaginal microbiome during pregnancy. Nat. Med. 25, 1001–1011.

49. Sommer, F., and Bäckhed, F. (2013). The gut microbiota--masters of host development and physiology. Nat. Rev. Microbiol. 11, 227–238.

50. The Human Microbiome Project Consortium. (2012). Structure, function and diversity of the healthy human microbiome. Nature 486, 207–214.

51. Walther-António, M.R.S., Chen, J., Multinu, F., Hokenstad, A., Distad, T.J., Cheek, E.H., Keeney, G.L., Creedon, D.J., Nelson, H., Mariani, A., et al. (2016). Potential contribution of the uterine microbiome in the development of endometrial cancer. Genome Med. 8, 122.

52. Wang, J., and Jia, H. (2016). Metagenome-wide association studies: fine-mining the microbiome. Nat. Rev. Microbiol. 14, 508–522.

53. Wang, J., Zheng, J., Shi, W., Du, N., Xu, X., Zhang, Y., Ji, P., Zhang, F., Jia, Z., Wang, Y., et al. (2018a). Dysbiosis of maternal and neonatal microbiota associated with gestational diabetes mellitus. Gut gutjnl-2018–315988.

54. Wang, P., Zheng, D., Wang, Y., and Liang, R. (2018b). One 3-oxoacyl-(acyl-Carrier-protein) reductase functions as 17 -hydroxysteroid dehydrogenase in the estrogen-degrading Pseudomonas putida SJ^β^TE-1. Biochem. Biophys. Res. Commun. 505, 910–916.

55. Zhang, W., Du, Y., Su, Z., Wang, C., Zeng, X., Zhang, R., Hong, X., Nie, C., Wu, J., Cao, H., et al. (2015a). IMonitor: A Robust Pipeline for TCR and BCR Repertoire Analysis. Genetics 201, 459–472.

56. Zhang, X., Zhang, D., Jia, H., Feng, Q., Wang, D., Liang, D. Di, Wu, X., Li, J.J.J., Tang, L., Li, Y.Y.Y., et al. (2015b). The oral and gut microbiomes are perturbed in rheumatoid arthritis and partly normalized after treatment. Nat. Med. 21, 895–905.

57. Morgan XC, Tickle TL, Sokol H, Gevers D, Devaney KL, Ward DV, Reyes JA, Shah SA, et al. (2012). Dysfunction of the intestinal microbiome in inflammatory bowel disease and treatment. Genome Biol. 13, R79–R79.

