## Supplementary Figures for "Life history recorded in the vagino-cervical microbiome"

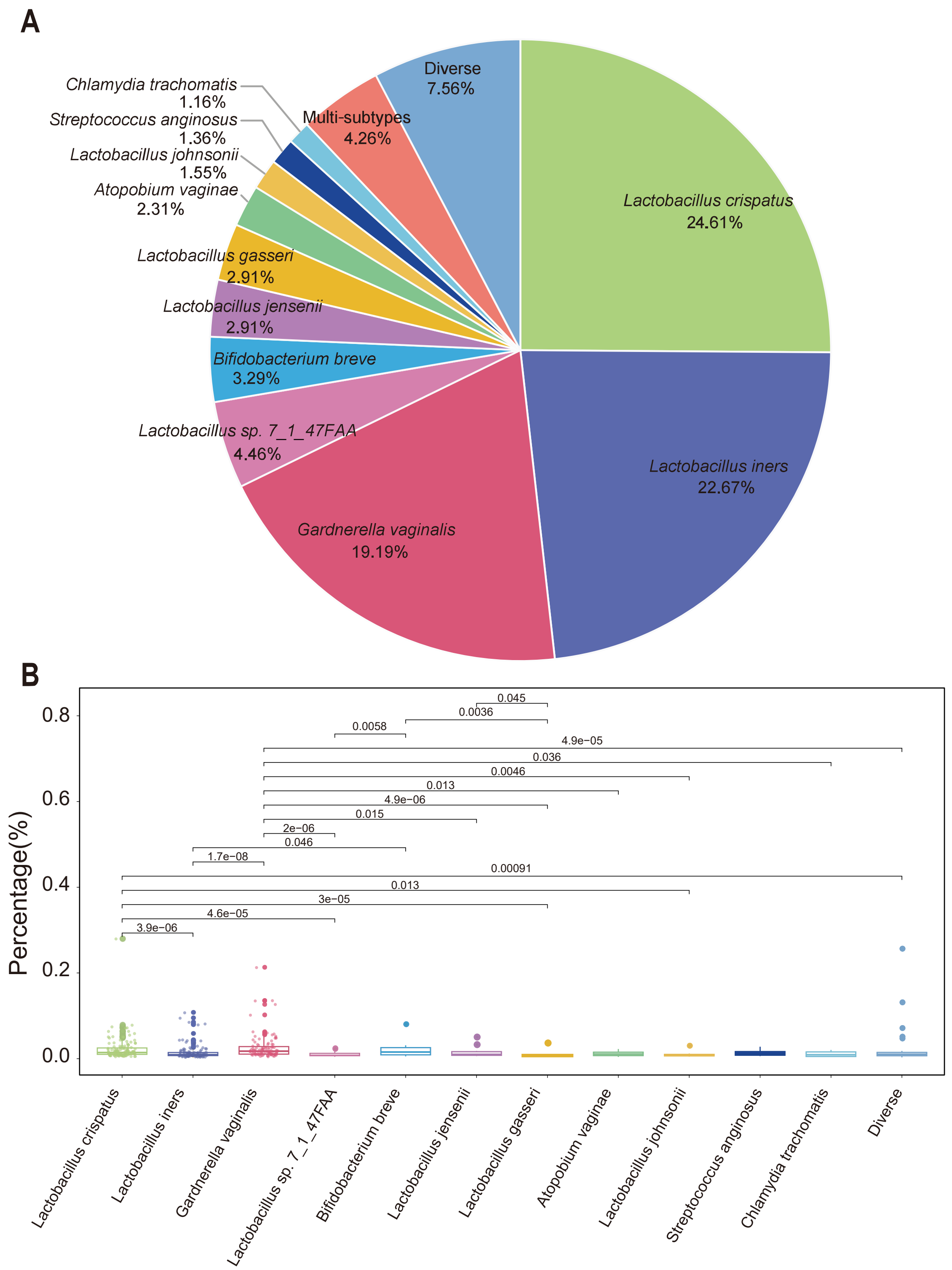


**Figure S1. Representative vaginal microbiota types identified in 516 women.**
(A) The ratio of different vaginal microbiota types. The species whose relative abundance account for more than 50% in an individual are selected as an identified type. Species all account less than 50% of the microbiota in an individual is identified as diverse type. The species types that represent less than 1% of individuals are labeled together as ‘Multi-types’.
(B) Percentage of non-human sequences in the dominate vaginal types (except Multi-types). Generalized linear model (GLM) is used to calculated the difference among the vaginal types. The boxes denote the interquartile range (IQR) between the first and third quartiles (25th and 75thpercentiles, respectively), and the line inside the boxes denote the median. The whiskers denote the lowest and highest values within 1.5 times the IQR from the first and third quartiles, respectively. Dots represent data points and the darker colored dots represent data points beyond the whiskers.

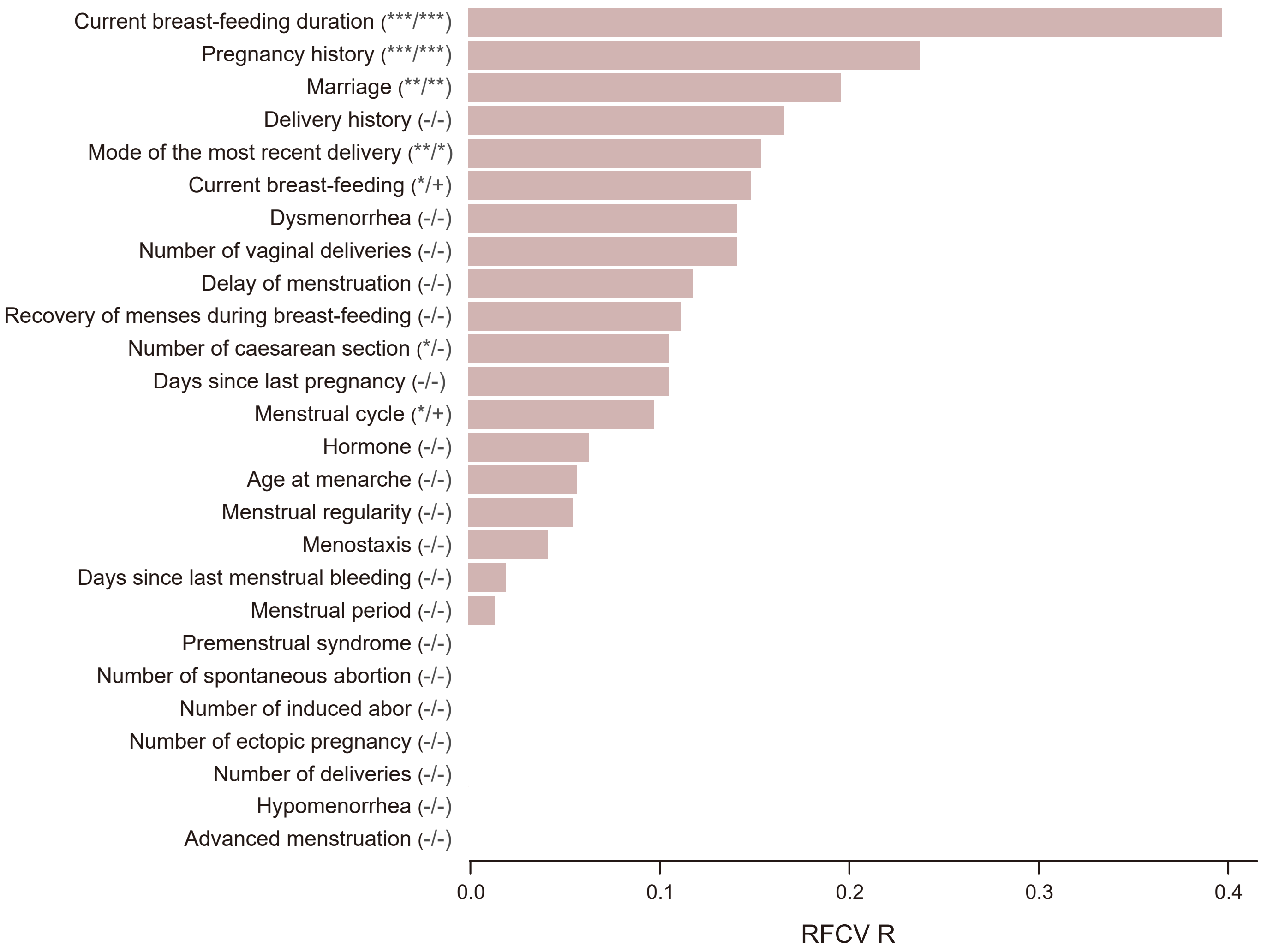


**Figure S2. Factors from female life history questionnaire influencing the vagino-cervical microbiome in the validation cohort.**

Female life history questionnaire entries on the vagino-cervical microbiome, ordered according to their 5-fold cross-validated random forest (RFCV) importance on the microbiome composition.

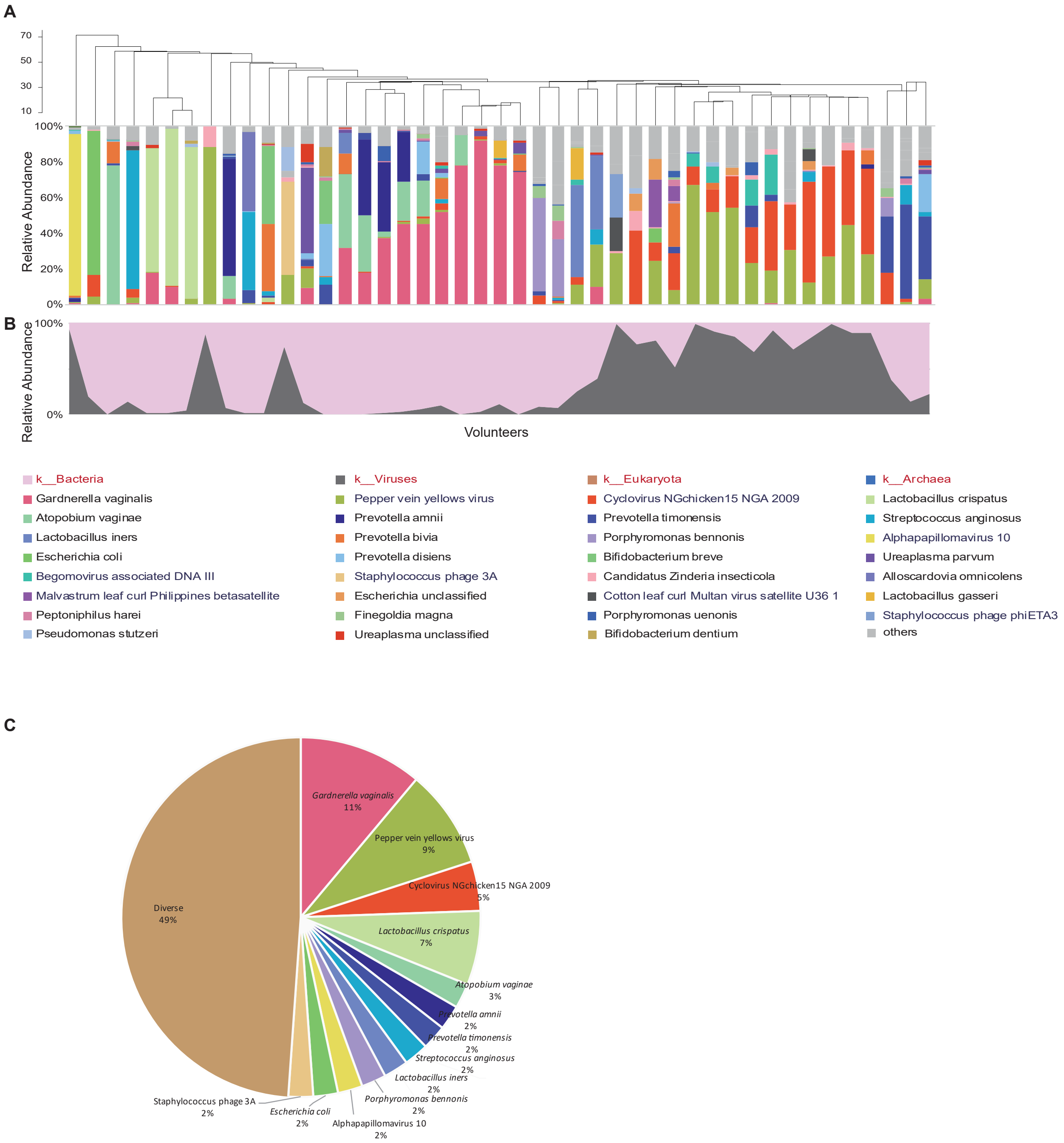


**Figure S3. Vagino-cervical microbiome of the postmenopausal women in validation cohort.**

The microbial composition in each sample at the species level (A) and kingdom level (B) according to MetaPhlAn2 is shown. The samples were hierarchically clustered (R base hcluster function with centroid linkage based on Euclidean distance. Taxa names in red, black and blue denote kingdom, bacteria and viruses, respectively. (C) The ratio of different vaginal-cervical microbiota types. The species whose relative abundance account for more than 50% in an individual are selected as an identified type. Species all account less than 50% of the microbiota in an individual is identified as diverse type.

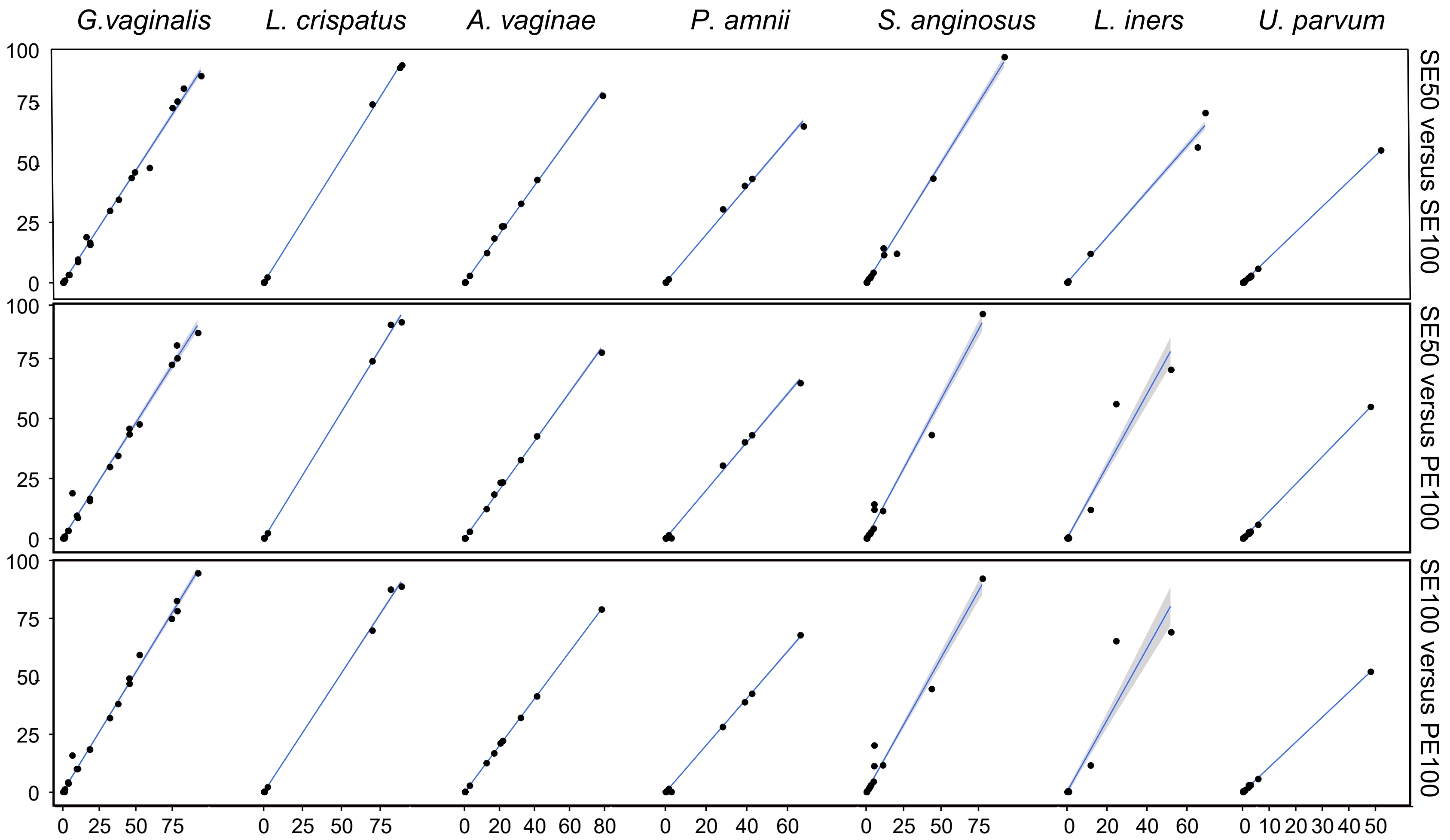


**Figure S4. Assessment reads length influence on abundance profile.**
For the 45 taxonomically more diverse postmenopausal samples in the validation set, we now include comparisons for the taxonomic profile from PE100, pseudo-SE100 and pseudo-SE50 by truncating the paired reads. The panel shows the top most abundance species in postmenopausal samples. Each point is a sample. X axis is the sample abundance from one of the PE100, pseudo-SE100, pseudo-SE50. And Y axis is the sample abundance from another mode. The fitting blue line is robust linear model regress one mode against another mode.

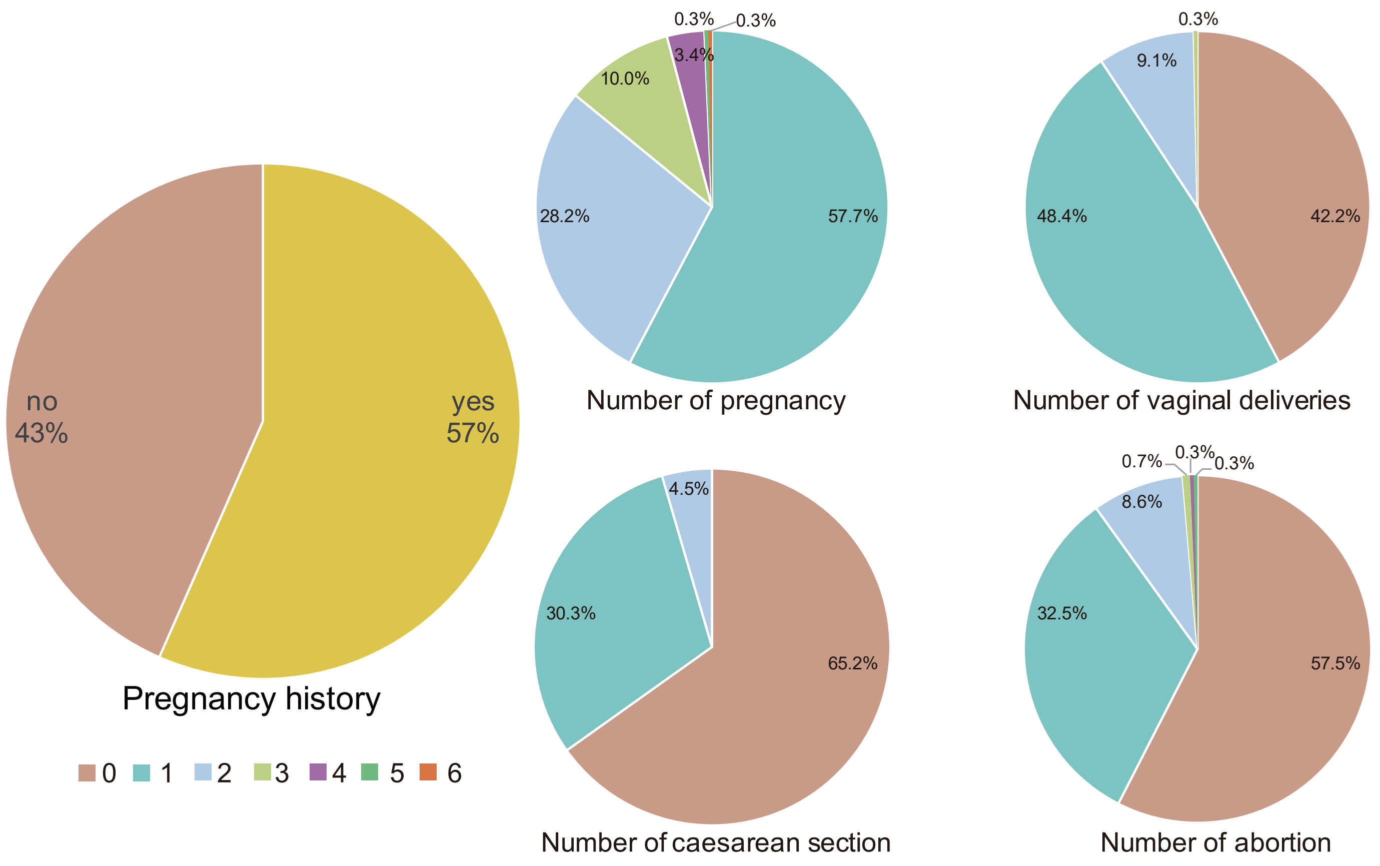


**Figure S5. The ratio of volunteers with breakdown of pregnancy history in the initial study cohort.**

The parts with different colors represent for 1-6 times of having the experience of pregnancy, delivery, caesarean section and abortion for one individual.

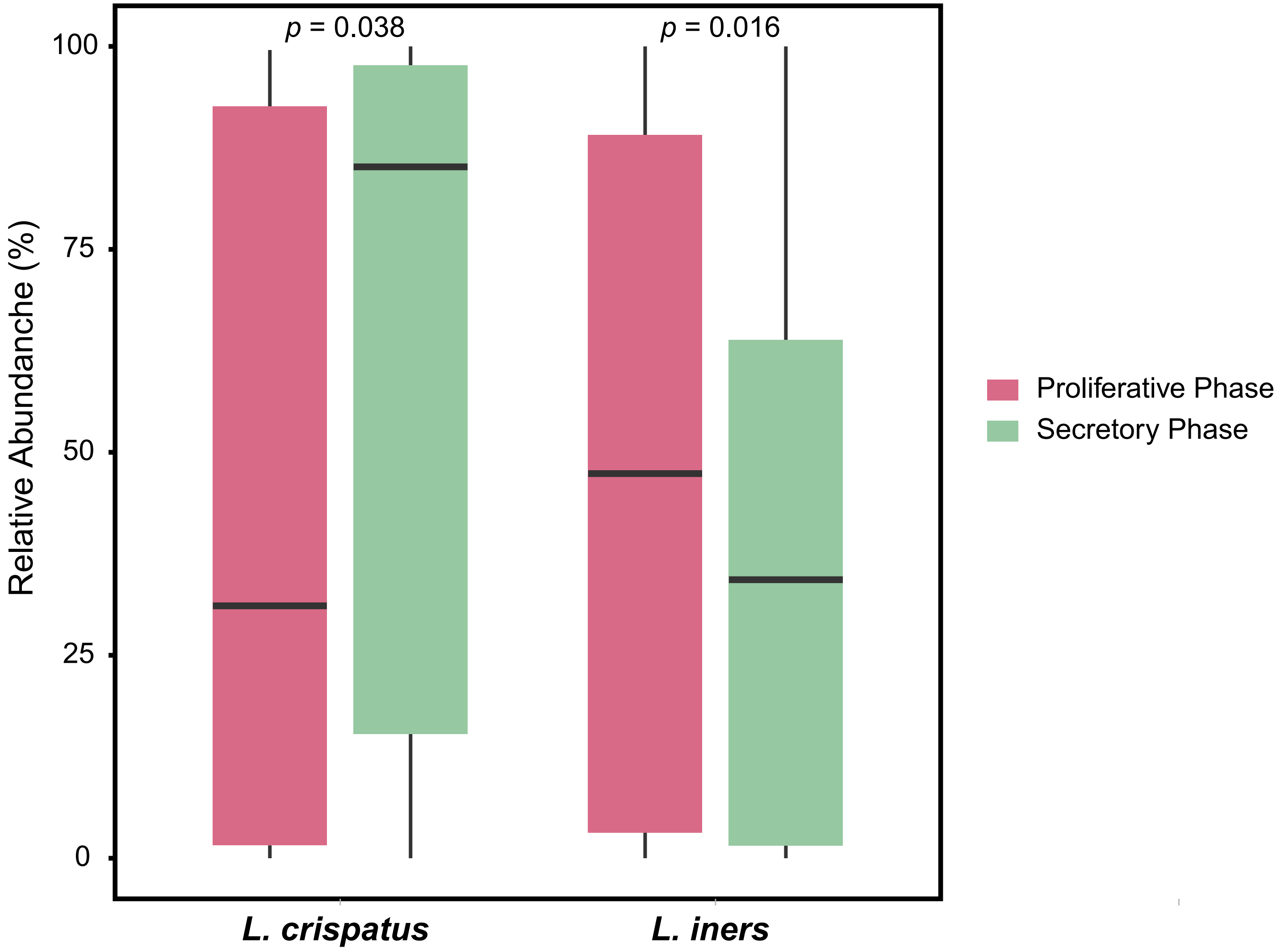


**Figure S6. The shift of *L. crispatus* and *L. iners* that shows signitificant difference during menstrual cycle in the initial study cohort** (Wilcoxon ranked sum test, $p<0.05$)**.** The length of menstrual cycle with each woman was normalized to 28 days. The proliferative phase is the end of period to 14th day, secretory phase is the 15th to 28th day of menstrual cycle. The boxes denote the interquartile range (IQR) between the first and third quartiles (25th and 75th percentiles, respectively), and the line inside the boxes denote the median.

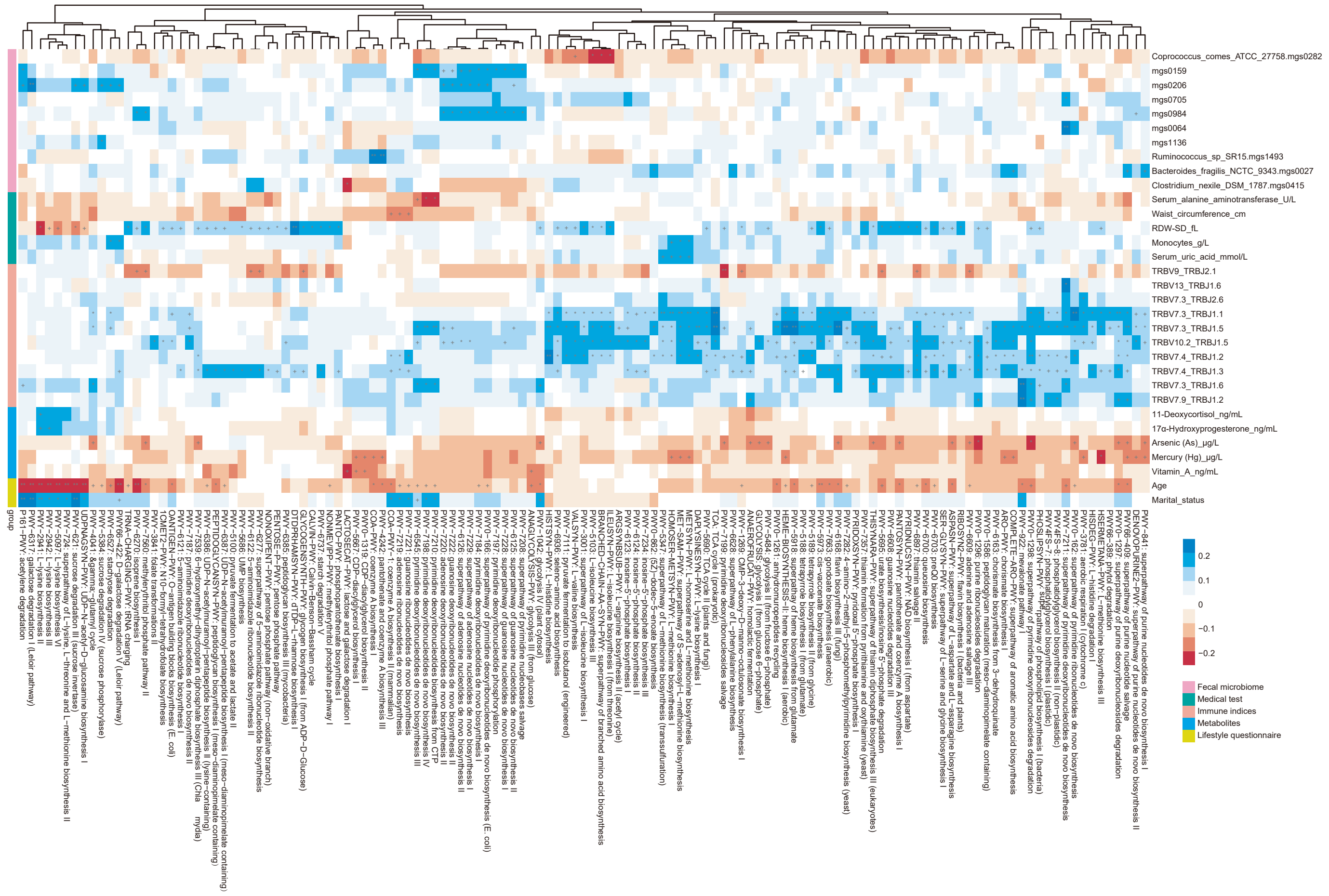


**Figure S7. Association between functional pathways of the vagino-cervical microbiome and other omics in the initial study cohort.**

Spearman's correlation between pathway profiles according to HUMAnN2 and the numerical data from other omics. Only those having significant associations with at least one pathway are shown (Q-value < 0.1, marked by an asterisk).

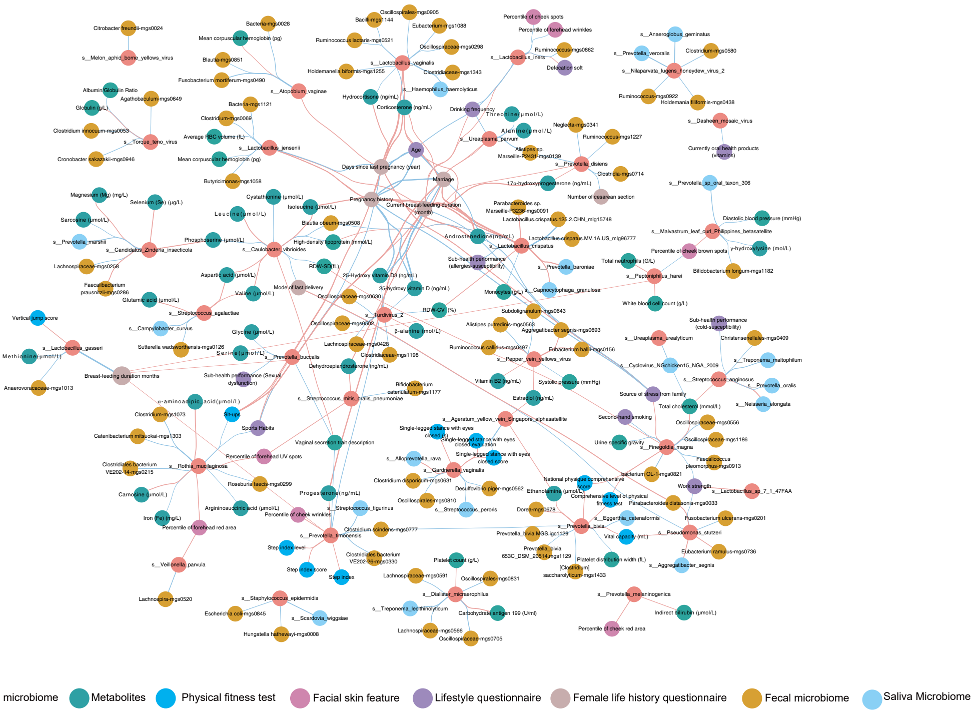


**Figure S8. Wisdom of the crowds for the association network between vagino-cervical microbial species and other omics data in the initial study cohort and the validation cohort.** Results from generalized linear model with penalty (cv. glmnet), random forest (RFCV) and Spearman's correlation are integrated and then visualized in CytoScape. The correlation is shown when the same enrichment direction in both cohorts and BH-adjusted P-value <0.1. Red lines, negative associations; cyan lines, positive associations( total 124528  associations, 336  for women life factor, 124528  for gut microbe, 1408  for clinical index, 4018  for metabolite, 1176  for PHY, 720  for PSYCH, 731 for SKIN, 1680  for lifestyle, 22876 for saliva microbe).
